# The transcription factor BCL11A defines a distinctive subset of dopamine neurons in the developing and adult midbrain

**DOI:** 10.1101/2020.10.06.327940

**Authors:** Marianna Tolve, Ayse Ulusoy, Khondker Ushna Sameen Islam, Gabriela O. Bodea, Ece Öztürk, Bianca Broske, Astrid Mentani, Antonia Wagener, Karen van Loo, Stefan Britsch, Pengtao Liu, Walid Khaled, Stephan Baader, Donato A. Di Monte, Sandra Blaess

**Affiliations:** Neurodevelopmental Genetics, Institute of Reconstructive Neurobiology, University of Bonn School of Medicine & University Hospital Bonn, Bonn, Germany; German Center for Neurodegenerative Diseases (DZNE), Bonn, Germany; Section for Translational Epilepsy Research, Department of Neuropathology, University of Bonn Medical Center, Bonn, Germany; Institute of Molecular and Cellular Anatomy, Ulm University, Ulm, Germany; School of Biomedical Sciences, Li Ka Shing Faculty of Medicine, The University of Hong Kong, China; Department of Pharmacology, University of Cambridge, Cambridge, UK; Institute of Anatomy, Anatomy and Cell Biology, University of Bonn, Bonn, Germany

## Abstract

Midbrain dopaminergic (mDA) neurons are diverse in their projection targets, impact on behavior and susceptibility to neurodegeneration. Little is known about the molecular mechanisms that establish this diversity in mDA neurons during development. We find that the transcription factor Bcl11a defines a subset of mDA neurons in the developing and adult murine brain. By combining intersectional labeling and viral-mediated tracing we show that Bcl11a-expressing mDA neurons form a highly specific subcircuit within the dopaminergic system. We demonstrate that Bcl11a-expressing mDA neurons in the substantia nigra (SN) are particularly vulnerable to neurodegeneration in an α-synuclein overexpression model of Parkinson’s disease. Inactivation of Bcl11a in developing mDA neurons results in anatomical changes, deficits in motor learning and a dramatic increase in the susceptibility to α-synuclein-induced degeneration in SN-mDA neurons. In summary, we identify an mDA subpopulation with highly distinctive characteristics defined by the expression of the transcription factor Bcl11a already during development.

## Introduction

Midbrain dopaminergic neurons (mDA) are anatomically organized into the substantia nigra, ventral tegmental area (VTA) and retro-rubral field (RRF). These anatomically defined areas contain subpopulations of mDA neurons that are characterized by distinct molecular profiles, distinct connectivity and distinct impacts on dopamine-modulated behavior (Engelhard et al., 2019; Poulin et al., 2018; Poulin et al., 2020). The SN consists of the pars compacta (SNc), pars lateralis (SNl) and pars reticularis (SNr). The majority of SN-mDA neurons is located in the SNc, a smaller population forms the SNl and only few mDA neurons are found in the SNr. SNl mDA neurons project to the tail of the striatum (TS) and have been shown to reinforce the avoidance of threatening stimuli (Menegas et al., 2018; Menegas et al., 2015). According to projection targets and functional output, the SNc is divided into a medial and lateral part. Medial SNc mDA neurons send their axons to the dorsomedial striatum (DMS) whereas projections of the lateral SNc target the dorsolateral striatum (DLS). Functionally, the firing of the DMS projecting mDA population may signal the valence of the outcome, while mDA neurons projecting to the DLS may signal a salience signal (appetitive or aversive) (Lerner et al., 2015). mDA neurons in the VTA project to the Nucleus accumbens (NAc), the olfactory tubercle (OT), prefrontal cortex (PFC) and amygdala. Similar to the SN, the VTA is anatomically divided into smaller domains, but here there is no clear relationship between the anatomical location of mDA cell bodies and the projection targets of VTA mDA neurons (Morales and Margolis, 2017). In addition to these anatomical mDA subgroups, mDA neurons have been defined based on their molecular profile. A comparative analysis of the currently available single cell gene expression studies led to the proposal that there are at least 7 molecularly-defined mDA subgroups. These do not necessarily correspond to mDA populations distinct by anatomical location, projection target and functional output (Poulin et al., 2020) and thus it is not clear to what extent these different levels of diversity can be reduced to a common denominator to define mDA diversity. Since anatomical position, molecular profile and connectivity are largely determined during development, a better understanding of the developmental factors that determine mDA subpopulations could deliver important new insights on how to define mDA subpopulations.

mDA neurons in the SNc degenerate in Parkinson’s disease (PD), leading to the cardinal motor symptoms of the disease. mDA neurons in the VTA are much less affected by the neurodegeneration in PD, but even within the SNc, neurons are not homogenous in their vulnerability: SNc-mDA neurons in the ventral tier appear to be more vulnerable than the ones in the dorsal tier. This selective vulnerability is found both in humans and in various PD models in rodents (Kordower et al., 2013). In mouse, the more vulnerable population has been shown to express the enzyme ALDH1A1 (Aldehyde Dehydrogenase 1 Family Member A1), while the less vulnerable subpopulation expresses the Calcium-binding protein Calbindin1 (CALB1) (Liu et al., 2014; Poulin et al., 2014). Since loss of ALDH1A1 function makes SNc-mDA neurons even more prone to neurodegeneration, ALDH1A1 seems unlikely to be the functionally decisive factor in the increased vulnerability (Liu et al., 2014). Nevertheless, these insights suggest that susceptibility to neurodegeneration in mDA subgroups could be genetically and developmentally pre-determined (Schwamborn, 2018).

Since transcription factors are key regulators of cell specification programs, we set out to identify transcription factors that are expressed in subpopulations of mDA neurons in the developing and adult brain. We discovered that the C2H2 zinc finger transcription factor BCL11A (B cell CLL/lymphoma) is expressed in a subpopulation of mDA neurons from embryogenesis to adulthood. BCL11A, also known as CTIP1 (Chicken ovalbumin upstream promoter transcription factor-interacting proteins 1), is a transcriptional repressor and a dedicated subunit of the mammalian SWI/SNF complex, a polymorphic assembly of at least 14 subunits (encoded by 28 genes) that functions as an ATP-dependent chromatin remodeler (Kadoch et al., 2013; Simon et al., 2020). In the mouse, BCL11A regulates neuronal fate determination during cortex and spinal cord development (Simon et al., 2020). In humans, BCL11A haploinsufficiency results in neurodevelopmental disorders characterized by developmental delay, mild to severe intellectual disability and behavioral problems (Basak et al., 2015; Deciphering Developmental Disorders Study, 2015; Dias et al., 2016; Peron et al., 2019). While these studies point to the importance of BCL11A in the development of the central nervous system (CNS), the function of BCL11A in development and maintenance of the dopaminergic system has not been examined.

We show here that the Bcl11a-expressing mDA neurons represent a previously uncharacterized subpopulation of mDA neurons, which does not correspond to any of the previously defined anatomical and molecular mDA subgroups but forms a highly distinct subcircuit within the dopaminergic system. Moreover, we demonstrate that Bcl11a appears to be required for the normal function of these mDA neurons, since conditional inactivation of Bcl11a in mDA neurons leads to deficits in motor learning in the conditional knock-out mice, although no significant loss of SN-mDA neurons or mDA projections could be detected in the mutant animals. Moreover, we find that *Bcl11a* expression characterizes a subpopulation of SNc neurons highly vulnerable to α-synuclein-induced neurodegeneration and that, within these neurons, *Bcl11a*-mediated transcription is likely to activate or modulate neuroprotective pathways. In summary, our data demonstrate that Bcl11a expression defines a subset of mDA neurons with highly specific projection targets and that loss of BCL11A interferes with the functional integrity and resilience to injury of these neurons.

## Results

### Bcl11a is expressed in a subset of mDA neurons in the SN, VTA and RRF

We initially identified *Bcl11a* as a potential mDA-subset marker based on the expression pattern available on the Allen Brain Atlas (Developing Mouse Brain). To examine whether Bcl11a is indeed expressed in a subset of mDA neurons, we analyzed the expression of *Bcl11a* mRNA and BCL11A protein in combination with tyrosine hydroxylase (TH), the rate limiting enzyme in dopamine synthesis. In the neonatal and adult brain, *Bcl11a* mRNA was expressed in a subset of SN, VTA, RRF and caudal linear nucleus (CLi) neurons. In the SN, *Bcl11a*-expressing mDA neurons were localized to the SNl and the medial and dorsal SNc. In the VTA, *Bcl11a*-positive neurons were found throughout the VTA (**Figure 1A-F**, data not shown). The distribution of neurons positive for BCL11A protein and TH was comparable to the distribution of *Bcl11a/*TH positive neurons in the neonatal brain and in the VTA of adult brains (**Figure 1G-L**), but we could not detect BCL11A protein in mDA neurons in the SNc and SNl in the P30 brain (**Figure 1J,L**). Since *Bcl11a* mRNA is expressed in SN-mDA neurons at P30, albeit at lower levels than in the VTA, this may indicate that SN-mDA express BCL11A protein at lower levels than VTA-mDA and that these low protein levels are not detected by the anti-BCL11A antibody used in our study.

**Figure 1.**
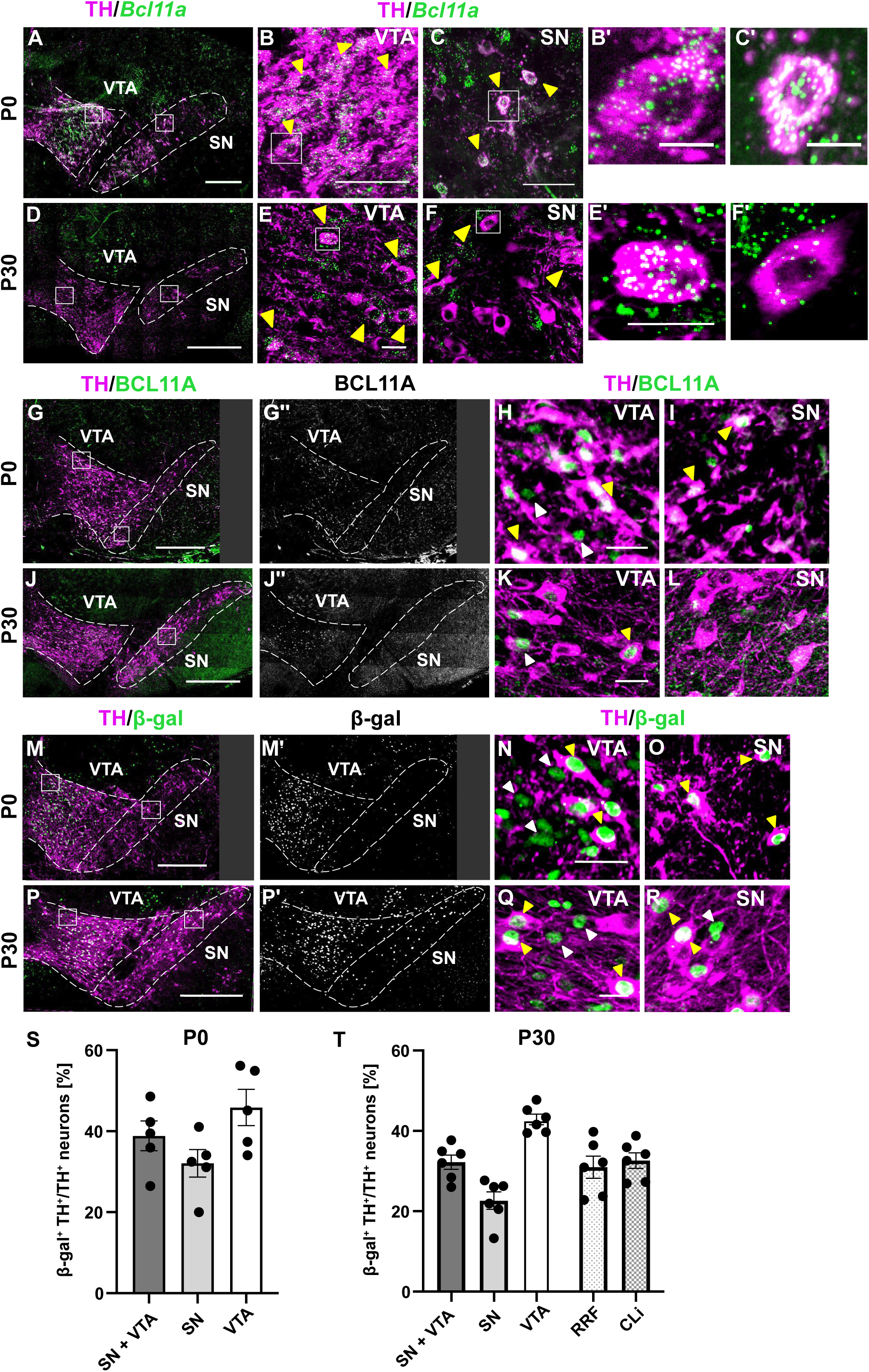
Bcl11a is expressed in a subset of SN, VTA, RRF and CLi neurons. **(A-F′)** Immunofluorescent staining for TH and RNAscope for *Bcl11a* mRNA on P0 **(A-C′)** and P30 **(D-F′)** coronal sections. (**B,C**) Higher magnification of the boxed areas in A; (**E,F**) Higher magnification of the boxed areas in D. **(B′,C′,E′,F′)** Higher magnification of the boxed area in B,C,E,F. Note that SN-mDA neurons express lower levels of *Bcl11a* mRNA than VTA-mDA neurons as evident by the density of fluorescent dots within the TH positive cell. **(G-L)** Immunofluorescent staining for BCL11A and TH on P0 **(G-I)** and P30 **(J-L)** coronal sections. **(H,I)** Higher magnification of the boxed area in G; **(K,L)** higher magnification of the boxed area in J. The immunostaining for BCL11A failed to detect the protein in the SN at P30. **(M-R)** Immunofluorescent staining for β-gal and TH on coronal sections of P0 **(M-O)** and P30 **(P-R)** *Bcl11a-lacZ* mice. **(N,O)** Higher magnification of the boxed area in M; **(Q,R)** higher magnification of the boxed area in P. Filled arrowheads indicate cells that are double positive for TH and BCL11A/*Bcl11a*/β-gal, empty arrowheads indicate cells that express BCL11A/*Bcl11a*/β-gal but are TH negative. **(S,T)** Percentage of TH-expressing neurons that are positive for β-gal in *Bcl11a-lacZ* mice at P0 (**S;** n=5 mice) and P30 (**T**; n=6 mice). Error bars indicate mean +/- SEM. Scale bars: 200 μm **(A,G-G′,M-M′)**, 500 μm **(D,J-J′,P-P′)**, 25 μm **(B,C,E,E′,F,F′,H,I,K,L,N,O,Q,R),** 10 μm **(B′,C′)**.

Next, we analyzed the distribution of *Bcl11a*-expressing cells using *Bcl11a^lacZ^* mice. In this mouse line, the *lacZ* allele is knocked into the endogenous *Bcl11a* locus and β-gal expression is restricted to cells that express *Bcl11a* (Dias et al., 2016). Indeed, the distribution of β-gal positive mDA neurons in neonatal and adult brain was comparable to the one of mDA neurons expressing *Bcl11a* mRNA (**Figure 1A-F, M-R, Supplemental Figure 1**). To test whether β-gal reliably marks BCL11A-expressing neurons, we performed double labeling for β-gal and BCL11A protein in neonatal and adult sections of the midbrain and found that the expression pattern of β-gal and BCL11A was largely overlapping in the VTA and in the cerebral cortex (**Supplemental Figure 2**). We then quantified the percentage of mDA neurons that express *Bcl11a* by counting β-gal and TH double-positive cells in *Bcl11a-lacZ* mice. β-gal was expressed in more than 40% of VTA-mDA neurons (neonatal: 45.84% +/- 4.467, adult: 42.84% +/- 1.326), in about one fourth of SN-mDA neurons (neonatal: 32.07% +/- 3.395, adult: 22.63% +/- 2.179) and in a third of RRF- and caudal linear nucleus (CLi)-mDA neurons (RRF: 30.97% +/- 2.757; CLi: 32.61% +/- 1.933, only analyzed in adult brain) (**Figure 1S,T**). Additionally, non-dopaminergic neurons expressing β-gal were found in the ventral midline of the midbrain, dorsal to the SN and within the VTA and SN area (**Supplemental Figure 1**).

To investigate the developmental time course of BCL11A expression in the ventral midbrain, we used immunostaining to analyze BCL11A protein expression between embryonic day (E)12.5 and E15.5. BCL11A was first expressed in the ventral midbrain at E12.5. At E12.5 and E13.5, BCL11A was mainly localized in the area just below the mDA progenitor domain and in a few differentiated TH-expressing mDA neurons. At E14.5 and E15.5, expression was found in a larger subset of mDA neurons, both in the forming SN and VTA (**Supplemental Figure 3**). These data demonstrate that Bcl11a-expressing mDA neurons constitute a subset of mDA neurons in the SN, VTA, RRF and CLi in the developing and adult brain.

### *Bcl11a*-expressing mDA neurons contribute to several known subpopulations of mDA neurons

A number of recent studies have described several subtype markers for mDA neurons, such as ALDH1A1, CALB1 and SOX6 (SRY-Box Transcription Factor 6). ALDH1A1 and CALB1 define complementary domains in the SNc (see introduction). In addition, ALDH1A1 is expressed in the ventral VTA while CALB1 shows a wide-spread expression in the VTA (Thompson et al., 2005; Wu et al., 2019). SOX6 expression is restricted to SN-mDA neurons and to neurons of the lateral VTA (Panman et al., 2014; Poulin et al., 2018). Since Bcl11a-expressing mDA neurons are broadly distributed within the mDA neuron-containing regions, we next asked whether the Bcl11a positive mDA population falls in one of these characterized subclasses. We used *Bcl11a-lacZ* mice for this analysis and performed triple immunostaining for β-gal, TH and the respective subset marker (ALDH1A1, CALB1 or SOX6) in the neonatal (**Figure 2A,B**) and adult brain (**Figure 2C,D**). Triple labeling for β-gal, TH and CALB1 or β-gal, TH and SOX6 at P0 showed that 24.33% +/- 3.810% of Bcl11a+ mDA neurons co-expressed CALB1 and 41.53% +/- 4.591% co-expressed SOX6 in the SN. In the VTA, 37.13% +/- 3.883% of mDA neurons co-expressed CALB1 and 14.09% +/- 1.771% co-expressed SOX6 at P0 (**Figure 2E,F**). At P30, triple labeling for β-gal, TH and CALB1 or ALDH1A1 showed that 45.18% +/- 5.903% of Bcl11a positive mDA neurons co-expressed CALB1 and 19.76% +/- 1.342% co-expressed ALDH1A1 in the SN, while 71.09% +/- 6.018% co-expressed CALB1 and 20.70% +/- 3.337% co-expressed ALDH1A1 in the VTA (**Figure 2G,H**). Thus, the *Bcl11a*- expressing mDA neurons do not clearly fall within one of these previously characterized mDA subpopulations. Of note, even though *Bcl11a*-expressing mDA neurons are mainly localized to the dorsal tier of the SNc and the SNl, which is typically characterized as CALB1-positive, only about half of them express CALB1 in the adult brain.

**Figure 2.**
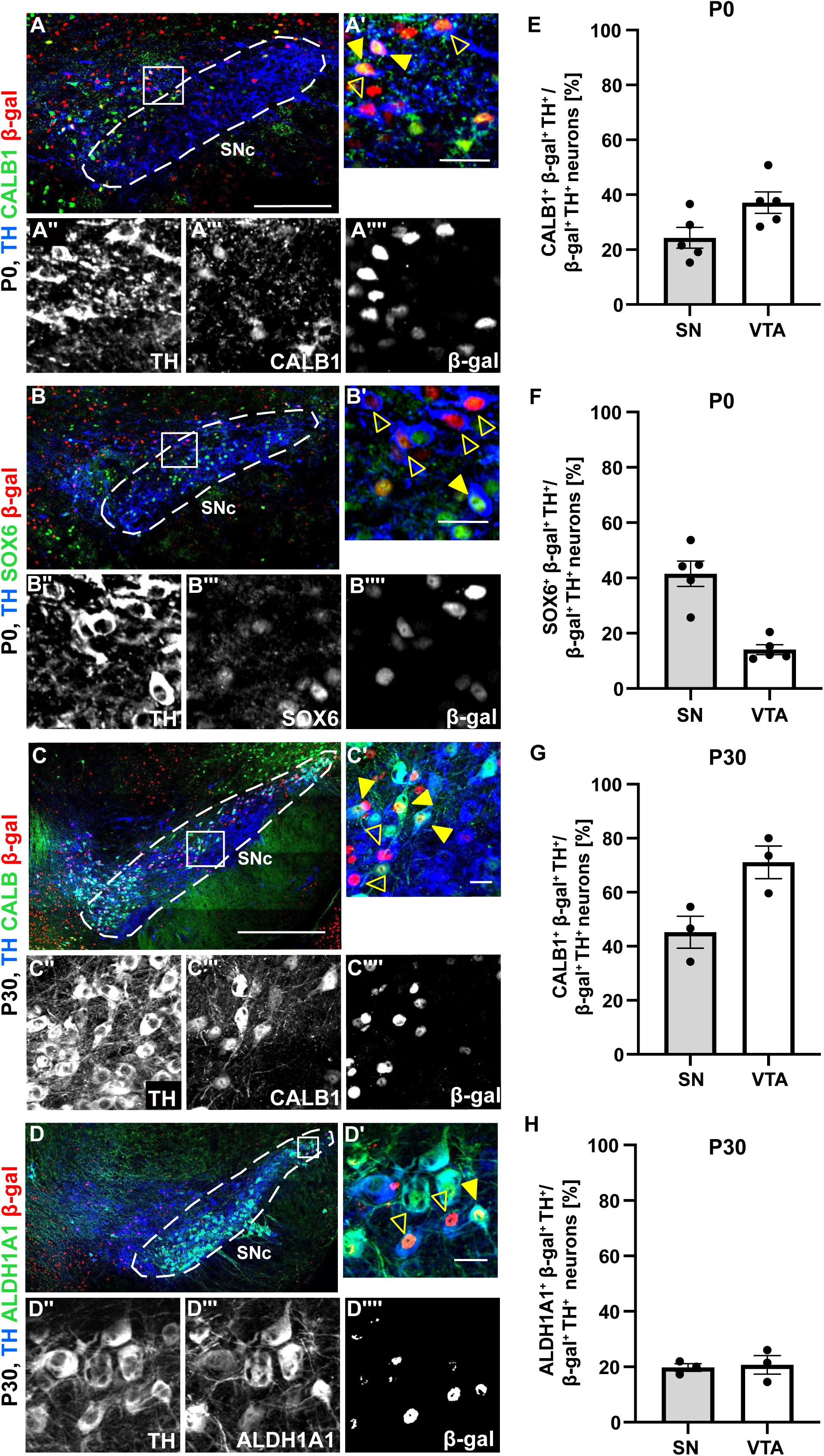
*Bcl11a*-expressing mDA neurons are a distinct mDA subset that does not correspond to previously defined mDA neuronal subpopulations. **(A,B)** Triple immunostaining for TH (blue), β-gal (red) and CALB1 (green) **(A)** or SOX6 (green) **(B)** in P0 *Bcl11a-lacZ* mice. **(A′-A′′′′,B′-B′′′′)** Higher magnification of the boxed area in A,B. **(C,D)** Triple immunostaining for TH (blue), β-gal (red) and CALB1 (green) **(C)** or ALDH1A1 (green) **(D)** in P30 *Bcl11a-lacZ* mice. **(C′-C′′′′,D′-D′′′′)** Higher magnification of the boxed area in C, D. Filled arrowheads indicate TH^+^ β-gal^+^ cells expressing the respective subset markers, unfilled arrowheads indicate TH^+^ β-gal^+^ cells negative for the respective subset marker. **(E-H)** Percentage of TH^+^ β-gal^+^ neurons that are positive for the respective subset marker at P0 (**E,F** n=5 mice) and at P30 (**G,H**, n=3 mice). Scale bars: 200 μm **(A,B)**, 500 μm **(C,D),** 25 μm **(A′-A′′′′,B′-B′′′′,C′-C′′′′,D′-D′′′′)**.

### *Bcl11a*-expressing mDA neurons form a subcircuit in the dopaminergic system

Next, we examined whether the Bcl11a-expressing subclass of mDA neurons contributes to specific subcircuits in the mDA system. To investigate projection targets of *Bcl11a*-expressing mDA neurons we used an intersectional genetic approach. This method combines a reporter allele or viral construct in which expression of a fluorescent reporter protein is driven by a tetracycline response element (TRE) in a Cre-dependent manner (Madisen et al., 2015; Poulin et al., 2018). To achieve specific activation of the reporter allele in mDA neurons we used a *Dat^tTA^* (tetracycline trans-activator driven by the *Dat* promoter) mouse line in conjunction with the *Bcl11a^CreER^* mouse line (Chen et al., 2015; Poulin et al., 2018) and the intersectional reporter mouse line Ai82D (Madisen et al., 2015) (**Figure 3A,B**). Since Bcl11a expression is already restricted to subsets of mDA neurons in the developing brain (**Supplemental Figure 3B-E**), CreER was activated during embryogenesis by administering Tamoxifen to pregnant females at E15.5. Distribution of the recombined (EGFP+) neurons in the adult brain was similar to the distribution of *Bcl11a*-mRNA expressing mDA neurons or β-gal-expressing cells in *Bcl11a-lacZ* mice at P0 and at P30 (**Supplemental Figure 4;** compare with **Figure 1** **and Supplemental Figure 1**). Analysis of the projection pattern established by the EGFP positive fibers showed that the recombined mDA neurons establish a highly specific innervation pattern in dopaminergic forebrain target areas. In target areas of VTA-mDA neurons (Morales and Margolis, 2017), *Bcl11a*-expressing mDA neurons strongly innervated the OT and the ventral and lateral shell of the NAc, but not the NAc core or the prefrontal cortex (**Figure 3C-E** and data not shown). Moreover, *Bcl11a*-expressing mDA neurons showed a highly specific projection pattern to the TS and the caudal DMS (**Figure 3C-F**). The TS is a known target of mDA neurons in the SNl, while the DMS is innervated by mDA neurons in the medial SNc (Lerner et al., 2015; Menegas et al., 2018; Poulin et al., 2018). Within these SN target areas, the densest innervation originating from recombined mDA neurons was observed in the ventral TS. Very sparse innervation was observed in the dorsolateral or rostral striatum.

**Figure 3.**
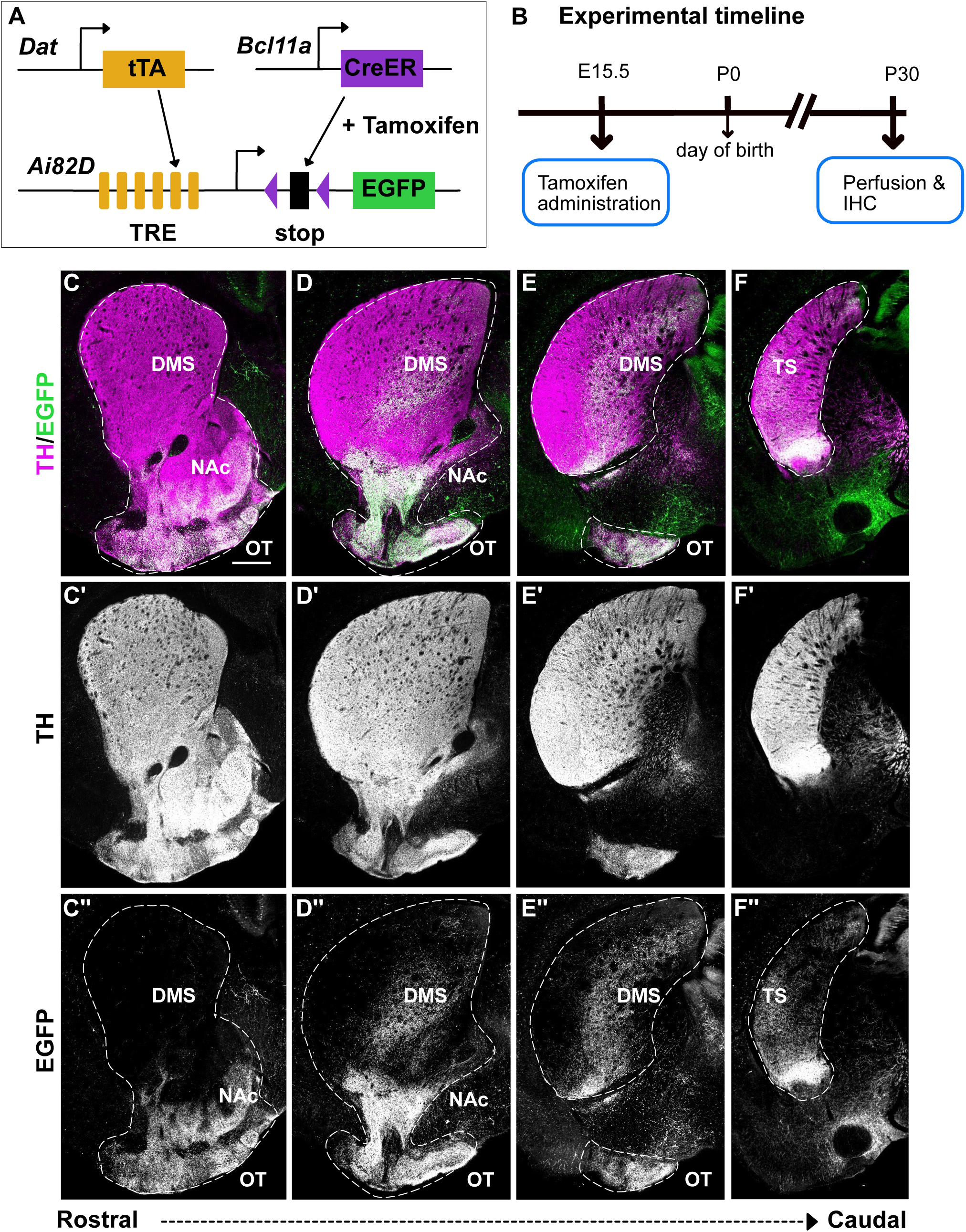
*Bcl11a*-expressing mDA neurons contribute projections to subcircuits of the dopaminergic system. **(A)** Schematic showing the intersectional fate mapping strategy to label *Bcl11a*-expressing mDA neurons and their projections. A *Bcl11a^CreER^* mouse line was used in combination with *Dat^tTA^* mice and an intersectional reporter mouse line (*Ai82D)*. EFGP is expressed only in cells positive for both CreER and tTa and only after CreER is activated by administration of Tamoxifen. **(B)** Schematic showing the experimental timeline. **(C-F′′)** Immunostaining for TH and EGFP in rostrocaudal levels of the striatal region showing that *Bcl11a*-expressing mDA neurons innervate specific subdomains of dopaminergic projection targets including the olfactory tubercle (OT, **C-E′′**), the Nucleus Accumbens (NAc)shell **(C-D′′)** the dorsomedial striatum (DMS, **C-E′′)** and the tail of the striatum (TS) **(F-F′′)**. Note that the most ventral part of the TS shows the highest density of EGFP positive fibers **(F-F′′)**. Mice analysed (n= 8 mice). Scale bar: 500 μm **(C-F′′)**.

In a next step, we investigated the projections of mDA neurons that express Bcl11a in the adult brain to (1) clarify whether *Bcl11a*-expression defines the same subset of mDA neurons in the embryonic and adult brain and (2) to clearly delineate the projection targets of *Bcl11a*-expressing neurons in the SN versus *Bcl11a*-expressing neurons in the VTA. To restrict the reporter activation to only Bcl11a-expressing neurons in either the SN or VTA, we introduced reporter constructs by injecting recombinant adeno associated viruses (rAAV) into the SN or VTA of adult animals. CreER was subsequently activated by the administration of Tamoxifen to achieve recombination (**Figure 4A,B; Supplemental Figure 5A,B**). Here we took both an intersectional approach, in which we introduced an intersectional reporter construct into *Bcl11a^CreER^*, *Dat^tTA^* mice (**Figure 4A,B**) and a non-intersectional approach in which we delivered a Cre-dependent reporter construct in *Bcl11a^CreER^* mice (**Supplemental Figure 5A,B**). Characterization of the VTA projection targets showed strong innervation of the OT and the ventral and lateral shell of the NAc (**Figure 4G-J, Supplemental Figure 5G-J**), consistent with the innervation pattern observed with the embryonic intersectional labeling (**Figure 3**). In some animals (n=7/10) in which VTA neurons were labeled more sparsely, the innervation pattern of *Bcl11a-expressing* VTA-mDA neurons in the OT showed a stripe-like pattern, indicating that axons from single neurons might innervate specific domains within the OT (**Supplemental Figure 5I**). Labeling of *Bcl11a*-expressing mDA neurons was scarcer in the SN compared to the VTA (**Figure 4C,D; Supplemental Figure 5C,D**), consistent with a lower percentage of *Bcl11a-*expressing mDA neurons in the SN than in the VTA (**Figure 1**; **Figure 5**). The sparse labeling allowed us to visualize patches of innervation likely derived from single SN axons (Brignani et al., 2020). These patches were restricted to the DMS and ventral TS, consistent with the findings obtained with the embryonically-induced intersectional labeling of *Bcl11a*-expressing neurons (**Figure 3**, **Figure 4E,F; Supplemental Figure 5E,F**). In conclusion, these results show that *Bcl11a* expression defines a subset of mDA neurons in embryogenesis and adulthood, which forms a highly specific subcircuit within the mDA system, despite a broad anatomical distribution of this subset within the mDA territory.

**Figure 4.**
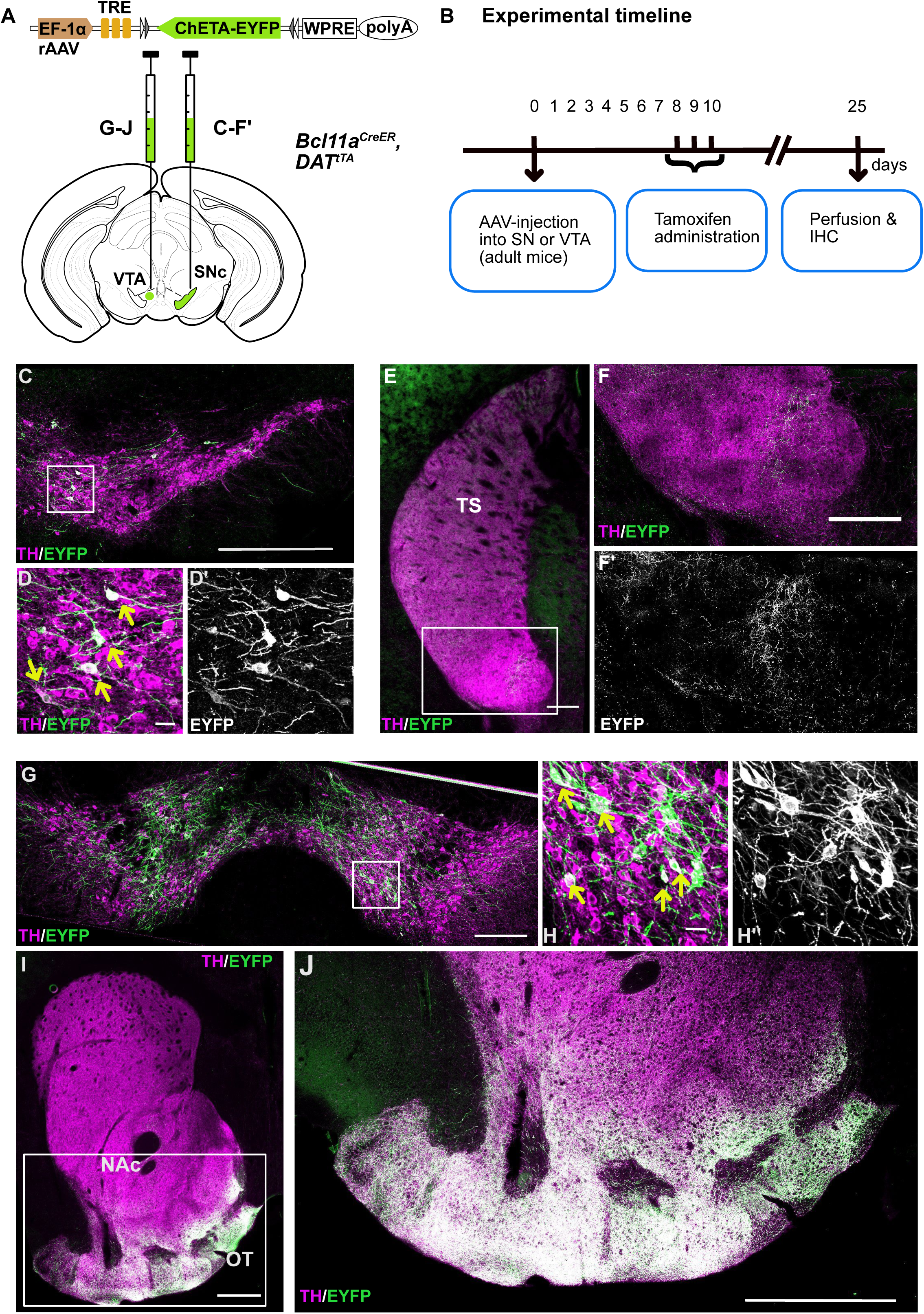
*Bcl11a*-expressing mDA neurons of the VTA and SN show a specific innervation pattern of forebrain targets**. (A)** Schematic showing injection of rAAV with an intersectional reporter construct into the SN or VTA of *Bcl11a^CreER^*^/+^; *Dat^tTA/+^* mice. **(B)**Schematic showing the experimental timeline. Tamoxifen administration 8 days after the virus injection results in expression of the reporter protein (EGFP) in *Bcl11a-*expressing mDA neurons. **(C-F′)** Immunostaining for TH and EGFP in the SN **(C-D′)** and the striatum **(E-F′). (E-F′)** EGFP^+^ fibers in the TS. **(F,F′)** Higher magnification of the boxed area in E. **(G-J)** Immunostaining for TH and the reporter protein in the VTA **(G-H′)** and NAc and OT **(I-J).** Yellow arrows indicate TH^+^ reporter protein^+^ neurons. Mice analyzed (n=2 mice for SN injections, n=3 mice for VTA injections). Scale bars: 500 μm **(C,G,I)**, 250 μm **(E,F,J),** 25 μm (**D,D′,H,H′**) .

**Figure 5.**
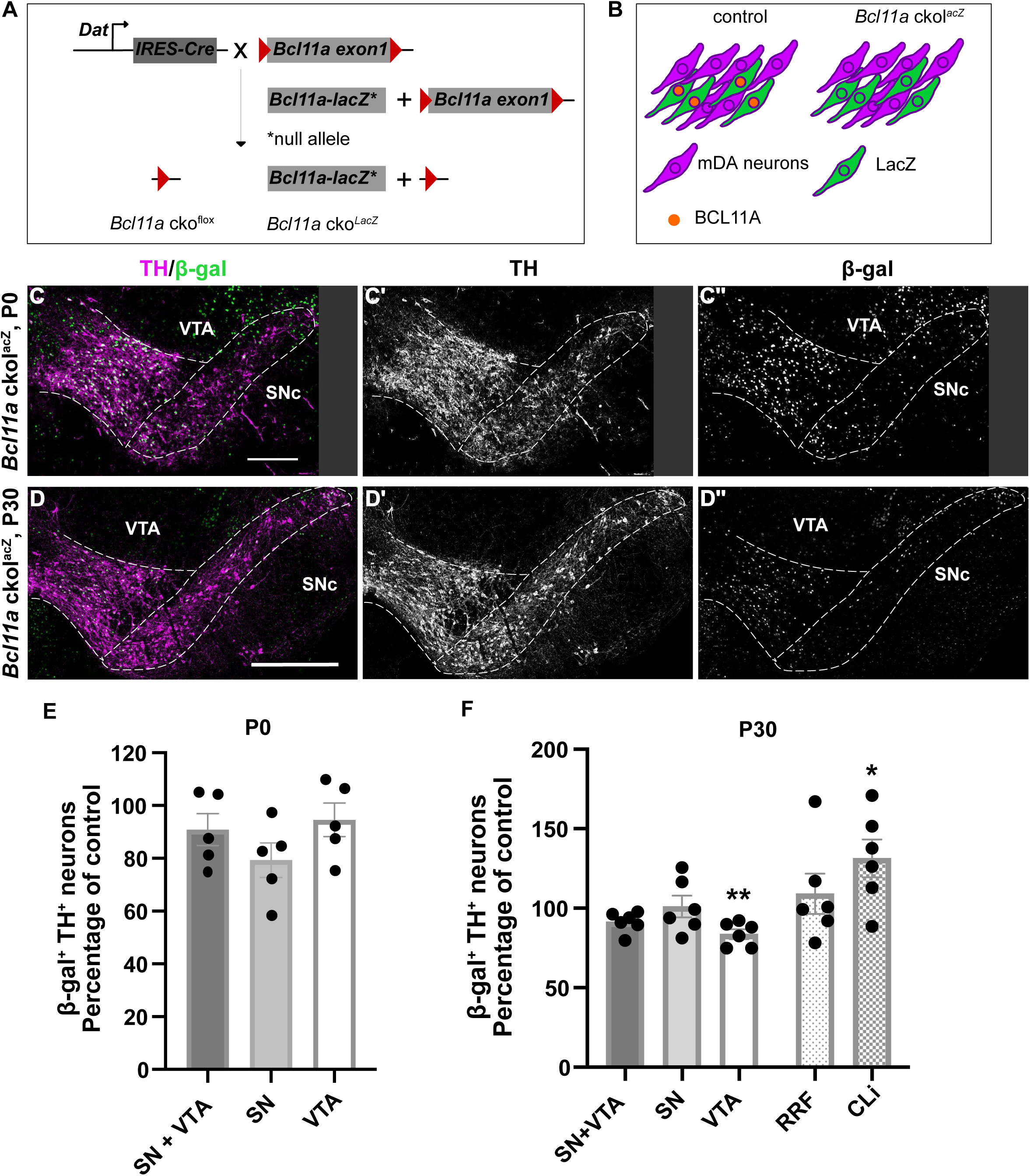
BCL11A is necessary for establishing the correct rostro-caudal position of *Bcl11a*-expressing mDA neurons in the VTA and CLi. **(A)** Conditional gene inactivation of *Bcl11a* in mDA neurons. *Bcl11a* cko mice were generated by crossing *Dat^IRES-Cre^* mice either with *Bcl11a^flox/flox^* mice (Genotype: *Dat^RES-Cre/+^, Bcl11a^flox/flox^*, termed *Bcl11a* cko^flox^) or with *Bcl11a^flox/lacZ^* mice (Genotype: *Dat^IRES-Cre/+^, Bcl11a^flox/lacZ^*, termed: *Bcl11a* cko^lacZ^). **(B)** In *Bcl11a* cko^lacZ^ mice, β-gal is a marker for Bcl11a-mDA neurons even after BCL11A expression is abolished. **(C-D′′)** Immunofluorescent staining for β-gal and TH on coronal sections of P0 **(C-C′′)** and P30 **(D-D′′)** *Bcl11a* cko^lacZ^ mice. **(E-F)** Quantification of TH^+^ β-gal^+^ neurons, expressed as percentage of control (*Bcl11a-lacZ* mice, control values are shown in Figure 1S,T). **(E)** There is no significant difference in the percentage of mDA neurons expressing β-gal in *Bcl11a* cko^lacZ^ mice (n=5) compared to *Bcl11a-lacZ* mice (n=5) at P0, indicating that Bcl11a-mDA neurons are established even in absence of BCL11A. **(F)** At P30, Bcl11a-mDA neurons are significantly decreased in the VTA and significantly increased in the CLi of *Bcl11a* cko^lacZ^ mice (n=6) as compared to *Bcl11a-lacZ* (n=6) indicating a shift in the distribution of Bcl11a-mDA neurons in absence of BCL11A **(F-G)**. Significance was determined by Welch’s t-test. \**p* < 0.05, ** *p* < 0.01. Error bars indicate mean +/- SEM. Scale bars: 200 μm **(C-C′′)** and 500 μm **(D-D′′’)**.

### Conditional gene inactivation of *Bcl11a* in mDA neurons results in a rostral-to-caudal shift of Bcl11a-mDA neurons from the VTA to the CLi

Since BCL11A is a transcription factor that has been shown to influence neuronal fate, neuronal morphology and migration (Simon et al., 2020), we investigated whether BCL11A is necessary for establishing and/or maintaining *Bcl11a*-expressing mDA neurons and their cell fate as well as their particular projection pattern. We generated a specific knock-out for *Bcl11a* in mDA neurons by crossing *Bcl11a^flox^* mice with *Dat^IRES-Cre^* mice (*Bcl11a* cko^flox^; Genotype: *Dat^IRES-Cre/+^, Bcl11a^flox/flox^*) (Bäckman et al., 2006; John et al., 2012). In a subset of mice, we introduced the *Bcl11a^lacZ^* allele (a null allele, (Dias et al., 2016)) into the conditional knock-out model (*Bcl11a* cko^lacZ^; Genotype: *Dat^IRES-Cre/+^, Bcl11a^flox/lacZ^*) (**Figure 5A**). In *Bcl11a* cko^lacZ^ mice, the *Bcl11a* expressing mDA population expresses β-gal even after inactivation of *Bcl11a*, giving us the possibility to analyze the effect of *Bcl11a* inactivation specifically in mDA neurons that would normally express BCL11A (termed: Bcl11a-mDA neurons) (**Figure 5B**). To confirm that Bcl11a expression was indeed absent in mDA neurons of *Bcl11a* cko^flox^ mice, we performed immunostaining and RNAScope experiments showing that BCL11A protein (**Supplemental Figure 6)** and *Bcl11a* mRNA (**Supplemental Figure 7)** were no longer expressed in mDA neurons. On the other hand, as expected, Bcl11a was still expressed in non-mDA neurons in the midbrain and the cerebral cortex (**Supplemental Figure 6 and 7**).

Despite the loss of BCL11A expression in mDA neurons, the anatomical organization of the mDA area in *Bcl11a* cko (both in *Bcl11a* cko^flox^ and *Bcl11a* cko^lacZ^) mice was comparable to controls (**Figure 5C, D** and data not shown). Quantification of the percentage of β-gal positive TH neurons in the SN and VTA in P0 *Bcl11a* cko^lacZ^ and *Bcl11-lacZ* mice showed no significant change in the number of Bcl11a-mDA neurons between mutant and control animals (**Figure 5E**). However, at P30, the percentage of β-gal positive TH neurons was significantly decreased in the VTA (average of three rostro-caudal levels corresponding to the levels in Supplemental Figure 1A,C,D) and significantly increased in the CLi (**Figure 5F**). When combining the numbers of β-gal positive TH neurons for the CLi level and the three VTA levels, no significant difference in the percentage of β-gal positive TH neurons could be detected between *Bcl11a* cko^lacZ^ and *Bcl11-lacZ* animals (**Supplemental Figure 8A**), indicating that the reduction of β-gal positive TH neurons in the VTA is likely due to a rostral-caudal shift in the position of Bcl11a-mDA neurons rather than a loss of cells. To examine this further, we compared the percentages of β-gal positive TH neurons at four rostral-caudal levels (three levels with VTA, one level with CLi). In addition to the significant increase in β-gal positive TH neurons in the CLi, we found that the percentage of β-gal positive TH neurons at the two rostral VTA levels (level 1 and 2) was significantly decreased in *Bcl11a* cko^lacZ^ compared to control mice (**Supplemental Figure 8B**). This resulted in a significant, systematic increase in the percentage of β-gal positive TH neurons from rostral-to-caudal in *Bcl11a* cko^lacZ^ mice, while there was a systematic decrease in β-gal positive TH neurons from rostral-to-caudal in control animals (**Supplemental Figure 8C**). The overall number of mDA neurons at the analyzed VTA and CLi levels was not significantly different between *Bcl11a* cko^lacZ^ and control mice (data not shown). No significant change in the percentage of β-gal positive TH neurons could be detected when comparing the rostral SN levels in control and *Bcl11a* cko^lacZ^ animals (data not shown), suggesting that the rostro-caudal shift of Bcl11a-mDA neurons in *Bcl11a* cko^lacZ^ animals is restricted to the VTA. Finally, stereological analysis of the total number of TH-positive mDA neurons or Nissl-stained neurons in the SNc of 12 months old *Bcl11a* cko and control mice did not show any evidence for cell loss in the *Bcl11a* cko mice (data not shown). These data indicate that the specific inactivation of *Bcl11a* in mDA neurons does not interfere with the generation or the survival of the Bcl11a-mDA neuronal population, but that it results in altered rostral-caudal positioning of Bcl11a-mDA neurons in the VTA.

If BCL11A is important for determining the localization of mDA neurons, it may also influence cell fate by regulating the expression of subset markers or the target specificity of Bcl11a-mDA projections. Triple labeling for β-gal, TH and CALB1 at P0 and P30, for β-gal, TH and SOX6 at P0 and for β-gal, TH and ALDH1A1 at P30 and quantification of triple labeled cells in *Bcl11a* cko^lacZ^ and control mice did not however reveal a significant change in the number of *Bcl11a*-mDA neurons co-expressing these markers in *Bcl11a* cko^lacZ^ mice as compared to *Bcl11a-lacZ* mice (**Figure 6**, compare with **Figure 2**). Next, we investigated whether inactivation of *Bcl11a* affects the targeting of projections arising from Bcl11a-mDA neurons. To this end, we examined the density of TH innervation in control and *Bcl11a* cko mice in the two areas with the highest innervation density from Bcl11a-mDA neurons, the ventral TS and the OT. In addition, we analyzed innervation density in the dorsal striatum and the dorsal TS. We found no significant difference in the density of the TH innervation in any of these areas when comparing control and *Bcl11a* cko mice (**Supplemental Figure 8D-J**). Taken together, these data suggest that the inactivation of *Bcl11a* and the resulting rostro-caudal shift in Bcl11a-mDA VTA neurons have no overt effect on the expression of known subset markers or the anatomical arrangement of Bcl11a-mDA projections.

**Figure 6.**
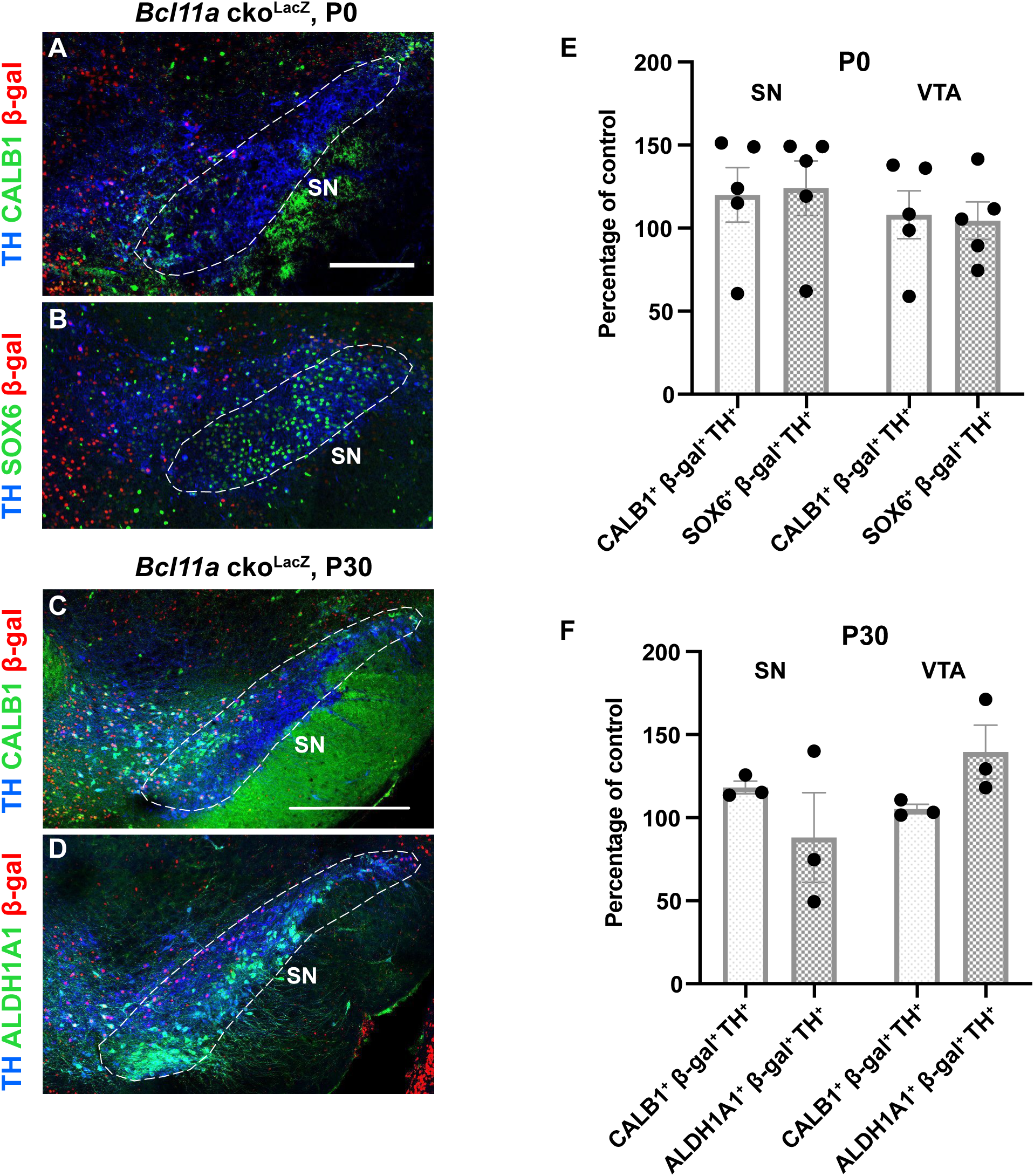
Expression of mDA subset markers in *Bcl11a*-expressing mDA neurons is not altered in absence of BCL11A. **(A,B)** Triple immunostaining for TH (blue), β-gal (red) and CALB1 (green) **(A)** or SOX6 (green) **(B)** in P0 *Bcl11a* cko^lacZ^ mice. **(C,D)** Triple immunostaining for TH (blue), β-gal (red) and CALB1 (green) **(C)** or ALDH1A1 (green) **(D)** in P30 *Bcl11a* cko^lacZ^ mice. **(E,F)** Quantification of triple positive neurons, expressed as percentage of control (*Bcl11a-lacZ* mice, control values are shown in Figure 2E-H). There is no significant difference in the percentage of TH^+^ β-gal^+^ mDA neurons expressing the analyzed subset markers in *Bcl11a* cko^lacZ^ mice compared to *Bcl11a-lacZ* mice at P0 or P30. N=5 cko and n=5 control mice were analyzed at P0, n=3 cko and n= 3 control mice were analyzed at P30. Significance was determined by Welch’s t-test. Error bars indicate mean +/- SEM. Scale bars: 200 μm **(A,B)**,; 500 μm **(E-F)**.

### Inactivation of Bcl11a in mDA neurons results in motor learning deficits

Inactivation of *Bcl11a* results in a rostral-to-caudal shift of the anatomical position of Bcl11a-mDA neurons in the VTA, but not in the SN. To investigate whether the loss of *Bcl11a* leads to functional impairment in the mDA system, either as a consequence of the subtle anatomical changes in the VTA or by a direct impact on mDA function, we examined a range of behaviors in *Bcl11a* cko and control mice. We chose tests to assess potential mDA neuronal functions associated with VTA- and/or SN. In the open field, distance moved was not altered in *Bcl11a* cko mice as compared to controls, indicating that spontaneous motor behavior was not affected in the mutant mice (**Figure 7A**). Decreased dopamine release from the VTA is associated with anxiety- and depressive-like behavior, which can be assessed by monitoring the activity in the center of an open field (Yacoubi et al., 2003; Tye et al., 2013). *Bcl11a* cko mice did not show a change in the frequency of crossing into the center and time spent in center or border area in the open field, indicating that the *Bcl11a* cko mice did not have an increased level of anxiety or depressive behavior **(Figure 7B,C**). Since inhibition of VTA-mDA neurons has been shown to lead to a decreased preference for social novelty (Bariselli et al., 2018), we used a social recognition test to examine the ability of the *Bcl11a* cko mice to distinguish familiar and unfamiliar mice. We did not find a significant alteration in the behavior of *Bcl11a* cko mice as compared to control mice (data not shown). Next, we focused on tasks in which dopamine release from SN-mDA neurons is thought to play a prominent role. Dopamine release in the striatum is crucial for voluntary movement and motor skill learning (Dodson et al., 2016; Wu et al., 2019). Motor coordination and balance were examined by monitoring the ability of mice to cross a balance beam (Luong et al., 2011) and we found no difference in the performance of *Bcl11a* cko mice and control mice in this task (**Figure 7D**). To examine whether motor skill learning is altered in *Bcl11a* cko mice, mice had to perform an accelerating rotarod test (Costa et al., 2004). In this task, control mice improved their performance over time, as reflected in a continuous increase in the time to fall over a 5-day training period. In contrast, *Bcl11a* cko mice did not show any improvement in their performance over time, indicating that they were not able to learn this motor task within the trial period (**Figure 7E**). These results show that inactivation of *Bcl11a* in mDA neurons results in defects in skilled motor learning suggesting that mDA neurons may be functionally impaired in the absence of *Bcl11a*.

**Figure 7.**
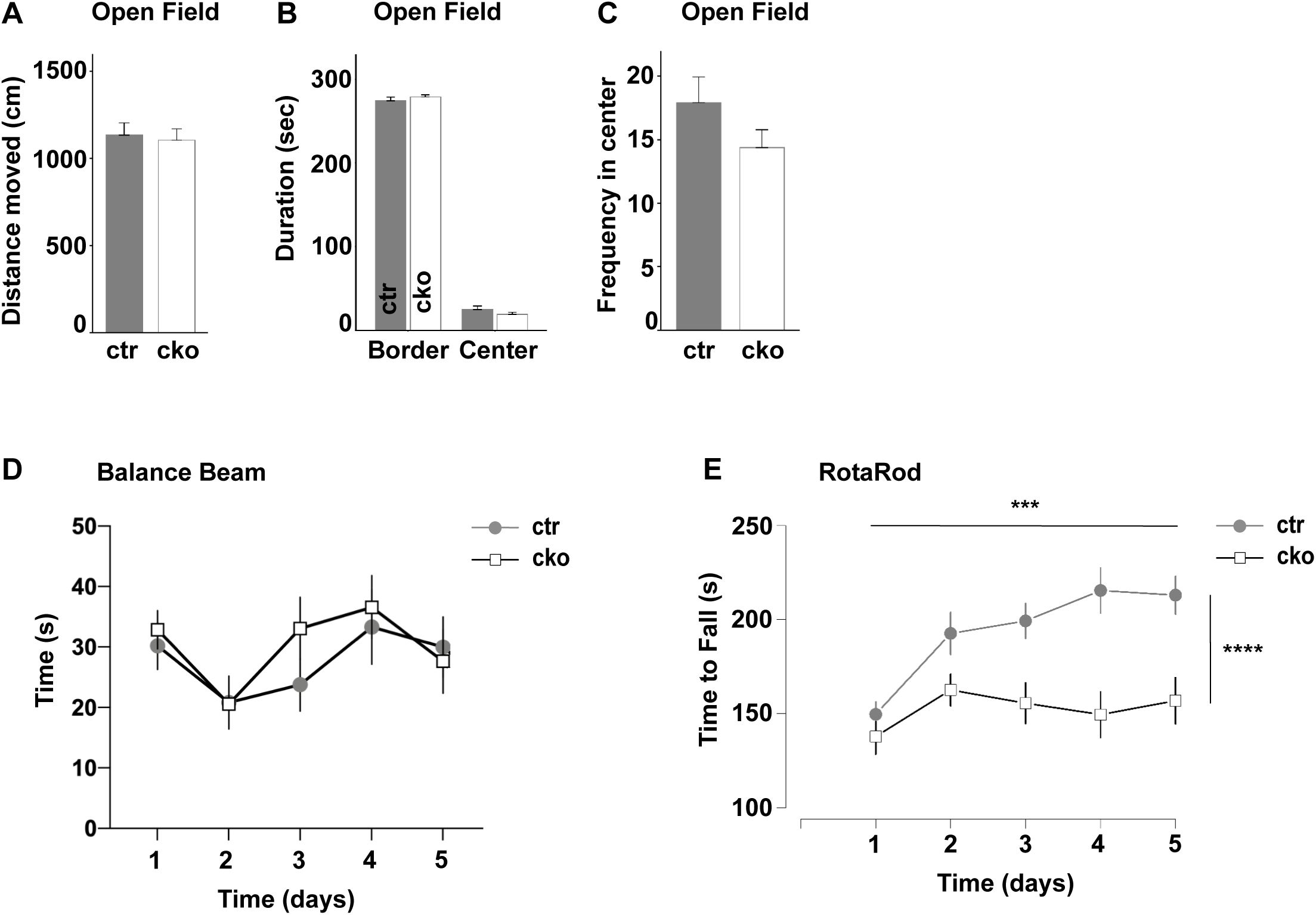
Inactivation of *Bcl11a* in mDA neurons results in specific defects in motor behavior. **(A-C)** Open field test (5-minute time window) revealed no significant difference between *Bcl11a* cko^flox^ (cko) and control mice (ctr) in distance moved **(A)**, in time spent in the center or border area **(B)** or the frequency of entering the center **(C)**. **(D**) Mice had to balance on a beam towards a food reward. The test was performed on 5 consecutive days. There was no significant difference between *Bcl11a* cko and control mice in their ability to cross a balance beam in any of the trials. **(E)** The rotarod test revealed that *Bcl11a* cko mice were not able to learn the motor task, since unlike in control mice the time to fall did not increase in *Bcl11a* cko mice within the 5 days of the trial period. *Bcl11a* cko (n= 12) and control mice (n= 13). Significance was determined by two-way ANOVA. \*\*\**p* < 0.001, **** *p* < 0.0001. p=7.226e-10 for genotype, p=0.0002017 for days and p=0.0076354 for interdependence between days and genotypes. Error bars indicate mean +/- SEM.

### Loss of Bcl11a enhances neuronal vulnerability to **α**-synuclein toxicity

The next set of experiments was aimed at elucidating the role of BCL11A expression in the context of neuronal challenges and neurodegenerative processes affecting mDA neurons in the SNc. The protein α**-**synuclein plays a key role in the pathogenesis of Parkinson’s and other human neurodegenerative diseases (Goedert et al., 2013). Its overexpression in animal models is associated with Parkinson-like pathology, including the degeneration of SNc-mDA neurons (Ulusoy and Di Monte, 2012). Here, adult control (n=5) and *Bcl11a* cko (n=5) mice were challenged with a single intraparenchymal injection of rAAVs carrying the DNA for human α**-**synuclein (**Figure 8A**). The unilateral injection targeted the right SNc where it caused robust overexpression of human α**-**synuclein within SNc-mDA neurons. No difference in overexpression was observed between control and *Bcl11a* cko mice after staining of SN-containing midbrain sections with a specific antibody against human α-synuclein (**Figure 8B**).

**Figure 8.**
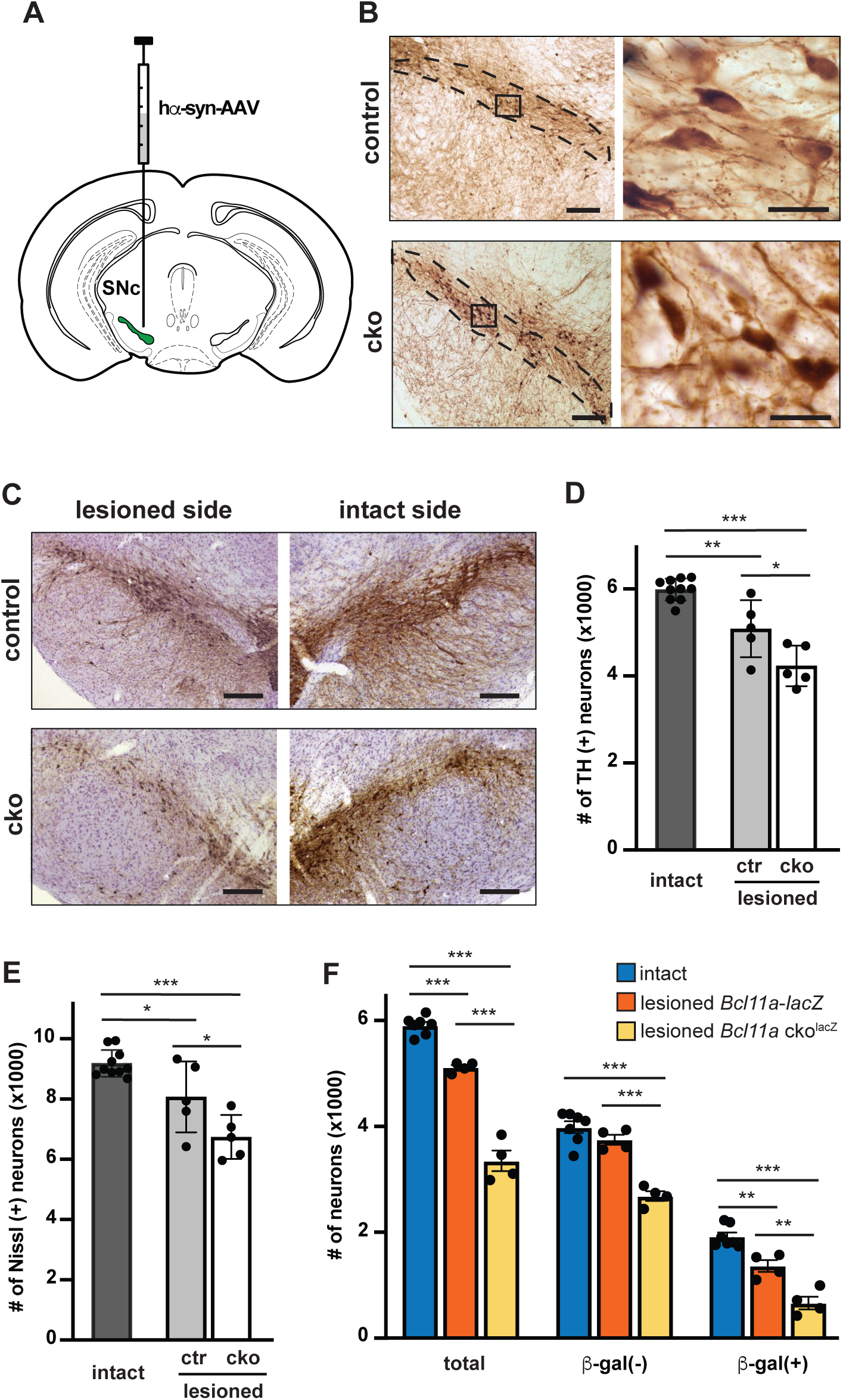
*Bcl11a* cko mice show increased neuronal vulnerability to α-synuclein toxicity. **(A)** Mice received a unilateral intraparenchymal injection of adeno-associated viral vectors (AAVs) carrying the DNA for human α-synuclein (hα-syn) into the SNc. Animals were sacrificed 8 weeks after this treatment. **(B)** Midbrain coronal sections comprising the SNc (delineated with dotted lines at low magnification) were immunostained with anti-human α**-**synuclein. Representative images from a control and a *Bcl11a* cko animal show neuronal expression of the transduced protein at low magnification (left panels). The right panels show a higher magnification of the boxed areas. **(C)** Midbrain sections were immunostained with anti-TH and counterstained with cresyl violet. Representative images from a control and a *Bcl11a* cko animal show TH-positive nigral neurons on right side of the brain lesioned with hα-syn-AAVs and on the contralateral intact side. **(D,E)** TH-immunoreactive (D) and Nissl-stained (E) neurons were counted stereologically in the intact (left) and lesioned (right) SNc of control (n=5) and *Bcl11a* cko (n=5) mice. Intact side values from control and cko animals were pooled together (dark gray bars). Light gray and empty bars show average values from the lesioned side of control (ctr) and cko mice, respectively. **(F)** Midbrain sections from *Bcl11a-lacZ* control (n=4) and *Bcl11a* cko^lacZ^ (n=4) mice were stained with anti-TH and anti-β-gal. Total neurons (β-gal negative and β-gal positive TH cells), β-gal(-) neurons (TH cells devoid of β-gal immunoreactivity) and β-gal(+) neurons (cells immunoreactive for both TH and β-gal) were counted using confocal stereology. Counts made in the intact SNc showed no difference between the two groups of animals. They were therefore pooled together (blue bars). Red and yellow bars show average values from the lesioned side of *Bcl11a-lacZ* and *Bcl11a* cko^lacZ^ mice, respectively. Significance was determined by one-way ANOVA followed by Tukey’s *post hoc* test for multiple comparisons. Error bars indicate mean +/- SEM. Scale bars: 100 μm (low magnification panels in **B; C**); 20 μm (higher magnification panels in **B**).

Midbrain sections were stained with a TH antibody, and immunoreactivity was compared in the left (intact side) and right (injected side) SNc of animals sacrificed 8 weeks after the rAAV injection. Labeling was overtly less robust in the lesioned right SNc of both control and *Bcl11a* cko mice. This loss of TH immunoreactivity appeared to be more pronounced, however, in sections from *Bcl11a* cko animals (**Figure 8C**). The number of TH^+^ neurons was then counted using a stereological method. Consistent with our previous quantification of Bcl11a-mDA neurons in *Bcl11a* cko and control animals (**Figure 5**), counts in the left (intact) SNc were similar between control and *Bcl11a* cko mice and were therefore averaged together as normal values (5,988 ± 79.4). Overexpression of α-synuclein caused a 15% reduction of TH^+^ cells in the lesioned right SNc of control mice, with cell counts averaging 5,086 ± 292.8. This loss was significantly more pronounced (4,231 ± 208.2 cells) in *Bcl11a* cko mice (**Figure 8D**). The possibility that the decrease in neuronal counts after rAAV injection may reflect a down-regulation of the phenotypic marker used for cell identification (i.e. TH) rather than actual cell degeneration was ruled out by counting the total number of Nissl-stained neurons. This number declined by 12% (from 9,185 ± 141.4 to 8,070 ± 525.6) and 27% (to 6,741 ± 326.1) in the lesioned SNc of control and *Bcl11a* cko mice, respectively (**Figure 8E**). These findings reveal that vulnerability to α-synuclein toxicity is markedly enhanced in the absence of *Bcl11a*, supporting the notion that *Bcl11a* expression is associated with transcription of genetic information involved in neuroprotective pathways.

Additional experiments using the same paradigm of rAAV-induced α-synuclein overexpression were carried out in *Bcl11a* cko^lacZ^ (n=4) and *Bcl11a*^flox/lacZ^ control (n=4) mice. Similar to the results in *Bcl11a* animals without the lacZ allele, α-synuclein toxicity caused a more severe loss of TH-immunoreactive neurons in the SNc of *Bcl11a* cko^lacZ^ mice (from 5,906 ± 64.4 to 3,349 ± 193.3 cells) as compared to *Bcl11a*^flox/lacZ^ controls (from 5,906 to 5,114 ± 47.9 cells) (**Figure 8F**, “total” values). In *Bcl11a*^flox/lacZ^ mice, confocal stereological counts were able to distinguish the effects of α-synuclein overexpression on Bcl11a-expressing SNc-mDA neurons (i.e. neurons that were both TH^+^ and β-gal^+^) *vs.* SNc-mDA devoid of Bcl11a (i.e. neurons that were TH^+^ but β-gal^−^). In these animals, counts of β-gal-positive and β-gal-negative mDA cells revealed that 29% (from 1,919 ± 78.3 to 1,365 ± 113.1 cells) of Bcl11a-expressing cells degenerated (**Figure 8F**, “β-gal(+)” values), whereas only 6% (from 3,986 ± 111.5 to 3,748 ± 95.1 cells) of β-gal-negative neurons were lost as a result of α**-**synuclein overexpression (**Figure 8F**, “β-gal(-)” values). These data indicate that *Bcl11a* expression characterizes a subpopulation of nigral DA neurons highly susceptible to α-synuclein-induced damage.

The number of β-gal-positive and β-gal-negative neurons was then counted in SNc of *Bcl11a* cko^lacZ^ animals. Results showed more dramatic toxic effects, since α**-**synuclein toxicity killed almost 65% (from 1,919 to 664 ± 117.0 cells) of β-gal-positive (**Figure 8F**, “β-gal(+)” values) and 33% (from 3,986 to 2,685 ± 91.4 cells) of β-gal-negative cells (**Figure 8F**, “β-gal(-)” values). Thus, in the absence of *Bcl11a* expression, β-gal-positive cells became even more vulnerable to neurodegeneration, consistent with a protective role of *Bcl11a*-mediated transcription. Interestingly, in *Bcl11a* cko^lacZ^ mice, a more severe neurodegenerative effect was observed not only on β-gal-positive but also β-gal-negative neurons. This latter finding suggests that inactivation of *Bcl11a* expression may have cell non-autonomous effects and result in widespread deleterious consequences for nigral tissue integrity.

## Discussion

### Expression of Bcl11a defines a previously uncharacterized subset of mDA neurons

mDA neurons are diverse in their gene expression, their anatomical location, their projection targets, their electrophysiological and functional properties (Farassat et al., 2019; Morales and Margolis, 2017; Poulin et al., 2020; Roeper, 2013). The link between anatomical position, gene expression, projection target and function has come more and more into focus in recent years. Several studies have shown that subsets of mDA neurons that share the expression of specific markers have also specific projection targets in the forebrain (Bimpisidis et al., 2019; Khan et al., 2017; Kramer et al., 2018; Poulin et al., 2018). Based on the currently available single-cell gene expression studies, it has been proposed that there are at least 7 molecularly defined mDA subgroups. Two of these subgroups are restricted to the SN, one encompasses cells in the SNc, RRF and lateral VTA, one is restricted to the linear nucleus and the remaining three are distributed in the medioventral VTA (Poulin et al., 2020). How and whether such subsets are already defined during development remains however unclear. Here we show that the transcription factor BCL11A defines a subpopulation of mDA neurons starting in embryogenesis and throughout adulthood. Bcl11a-mDA neurons are widely distributed within the anatomically defined mDA nuclei, comprising about 40% of the VTA-mDA neurons and about 25% of the SN-mDA neurons. The Bcl11a-mDA population does not clearly fall into one of the molecularly defined populations described above (Poulin et al., 2020). Strikingly, despite the broad distribution of Bcl11a-mDA neurons across different anatomical and molecularly defined mDA subpopulations, Bcl11a-mDA neurons establish a highly selective innervation pattern within the mDA projection targets.

### Bcl11a-mDA neurons form a highly specific subset in the midbrain dopaminergic system

Dopaminergic subcircuits may encode a precise behavioral output by targeting a set of substructures within the classical dopaminergic projection targets and a number of studies have shown that specific subcircuits within the mDA system regulate very specific aspects of behavior (Engelhard et al., 2019; Heymann et al., 2020; Menegas et al., 2018). Whether such behavioral modules are consistent with genetically determined populations is just starting to be examined (Heymann et al., 2020).

Bcl11a-mDA of the VTA project to the medial and ventral shell of the NAc and the OT while Bcl11a-mDA of the SN show a highly selective innervation of the ventral TS and the caudal DMS. This specific innervation pattern suggests that this genetically defined subpopulation may modulate a specific subset of dopamine-influenced behaviors.

From an evolutionary standpoint, the mDA system may consist of different modules with different functions that were built on top of a basic DA system mainly concerned with basic behavior just as food-seeking and reward behavior (Schultz, 2019). If there are such modules one would expect that they are genetically defined by the expression of specific transcription factors or sets of transcription factors. Bcl11a could potentially define such a behavioral-anatomical module and it will be interesting to investigate in the future whether mDA neurons that are characterized by *Bcl11a*-expression in the SN and VTA form such a functional module within the mDA system.

### Function of BCL11A in developing and mature mDA neurons

BCL11A is a zinc finger transcription factor that acts mainly as a repressor but it is also part of an ATP dependent chromatin remodeling complex in neural tissue. *Bcl11a* is expressed in many types of neurons in almost every region of the CNS (Allen Brain Atlas), but its molecular function has only been studied in a few regions so far (Simon et al., 2020). In the cortex, BCL11A is important for the specification of cortical neurons that project to subcerebral areas (Canovas et al., 2015; Woodworth et al., 2016) and it controls the acquisition of sensory area identity and the establishment of sensory input fields (Greig et al., 2016). Moreover, BCL11A regulates the migration of cortical projecting neurons (Wiegreffe et al., 2015). In the dorsal spinal cord, BCL11A is required for neuronal morphogenesis and sensory circuit formation (John et al., 2012). Whether it functions purely as a transcriptional repressor or also as part of the chromatin remodeling complex in these neurons remains to be investigated. In mDA neurons, the inactivation of *Bcl11a* resulted in a rostral-to-caudal shift of Bcl11a-mDA cell bodies, suggesting that BCL11A might play a role in regulating the expression of factors that control the migration of VTA-mDA neurons. In the cerebral cortex, BCL11A regulates cortical neuron migration by controlling the expression of the cell adhesion molecule Semaphorin 3c (Wiegreffe et al., 2015), but Semaphorin 3c is not expressed in mDA neurons (Kolk et al., 2009). Thus, it will be of great interest to investigate which other molecules are regulated by BCL11A in developing mDA neurons, in particular because little is known about the mechanisms underlying the migration of VTA-mDA neurons (Brignani and Pasterkamp, 2017). While the changes in cell body positioning appear to be restricted to the VTA, the loss of BCL11A results in SN-mDA neurons, which are functionally compromised and are more susceptible to neurodegeneration in the absence of BCL11A as evident from the results of the behavioral analysis and the α-synuclein overexpression experiments. The deficits in SN-mDA neurons demonstrate a novel role for BCL11A in regulating neuronal function and vulnerability that appears to be independent of its previously characterized role in cell fate specification.

Whether the effect of BCL11A on SN-mDA neurons is a consequence of BCL11A acting during the development of mDA neurons by altering their molecular and functional profile and/or whether BCL11A regulates the function and vulnerability of mDA neurons acutely in the adult brain cannot be discerned from the current study. Given that BCL11A appears to be continuously expressed in the same mDA subpopulation starting soon after mDA differentiation and throughout adulthood, a role of BCL11A during mDA development seems likely. However, only the analysis of a mouse model, in which *Bcl11a* is inactivated in mDA neurons during adulthood, will address this point conclusively.

In humans, pathogenic variants of BCL11A (mostly *de novo* mutations) result in neurodevelopmental disorders that have recently been classified as BCL11A-related intellectual disability (Peron et al., 2019). This syndrome is characterized by general developmental delay, microcephaly, speech delay and behavioral problems as well as a number of non-CNS related phenotypes. In some affected individuals, seizures or autism spectrum disorder have been reported (Peron et al., 2019). Given the widespread expression of BCL11A in the brain, it is unclear if any of these neurological symptoms in patients are associated with functional deficits in the dopaminergic system. Moreover, there are no reports of neurodegenerative phenotypes, but long-term observations of patients or case studies in adult or aged patients are not available, due to the limited number of cases reported and the bias towards diagnosis in childhood (Peron et al., 2019).

### Consequences of Bcl11a-inactivation in mDA neurons on behavior

*Bcl11a* cko mice show a defect in the learning of skilled motor behavior while spontaneous motor behavior or motor coordination is not affected. Since *Bcl11a* is specifically inactivated in mDA neurons in our mouse model, this particular behavioral phenotype must be caused by functional changes in mDA neurons (rather than by deficits in the cerebellum or motor cortex (Hikosaka et al., 2002; Li et al., 2017). Due to the highly specific innervation pattern of Bcl11a-mDA neurons in the forebrain and the fact that Bcl11a-mDA neurons comprise only a subset of mDA neurons, we were not able to address the nature of these functional changes, but we assume that they ultimately result in altered dopamine release in the target areas of Bcl11a-mDA neurons and that these alterations are severe enough to elicit a behavioral phenotype. Learning of motor skills is thought to be mediated by dopamine release in the dorsal striatum. It has been proposed that the plasticity in striatal medium spiny neurons that underlies initial skill learning during goal-directed actions requires dopamine release in the DMS, while dopamine release in the dorsolateral striatum (DLS) is important for optimal skill learning (the learning of sequential actions until a point is reached at which they can be performed with little effort or attention (Durieux et al., 2012; Graybiel, 2008). Based on this model and our observation that Bcl11a-mDA neurons in the SNc project to the DMS but not the DLS, the deficit in motor learning should have its origin in the inability of the mice to initiate the learning of skilled motor behavior. Interestingly, the ablation of *Aldh1a1*-expressing mDA neurons leads to an impairment in motor skill learning on the rotarod that is similar to the one observed in *Bcl11a* cko mice even though *Aldh1a1*-expressing neurons project almost exclusively to the DLS (Wu et al., 2019). Thus, it is possible that reduced dopamine release in both DMS and DLS and thus defects in both phases of learning lead to a similar overall defect in the acquisition of skill learning.

Finally, we cannot completely exclude that impaired dopamine release from Bcl11a-mDA neurons in the VTA may play a role in the inability of the *Bcl11a* cko mice to perform on the rotarod since they might be less motivated to perform the task. However, the mice did not show any deficits in any of the other tasks (balance beam, social recognition, open field) suggesting that a deficit in motivated behavior is unlikely to be the cause of the inability to learn the rotarod task.

### Vulnerability of Bcl11a neurons to degeneration

The mDA neurons in the ventral tier of the SNc is the one most vulnerable to neurodegeneration in Parkinson’s disease and animal models of the disease (Kordower et al., 2013). Based on studies in rodents, this population appears to coincide with ALDH1A1-expressing mDA neurons (Poulin et al., 2014). Here we demonstrate that BCL11A neurons are highly susceptible to neurodegeneration in an α-synuclein overexpression model, even though only a small percentage overlaps with ALDH1A1-expressing mDA neurons. Our data suggest that BCL11A serves as a marker of highly vulnerable mDA neurons, which are located primarily in the dorsal tier of the SNc and project to the caudal DMS. The DMS corresponds roughly to the caudate nucleus in the human brain (Burton et al., 2015). Dopaminergic deficits within the striatum are unevenly distributed in Parkinson’s disease patients and in general, neuroimaging studies show that the posterior putamen (corresponding to the DLS in rodents) has a more severe dopamine dysfunction than the caudate nucleus (Kish et al., 1988). This gradient of dopaminergic deficiency has been reported to be already apparent at early disease stages and to be largely maintained over the course of the disease (Nandhagopal et al., 2009). Nevertheless, some of the early symptoms associated with Parkinson’s disease are assumed to be based on functional deficits of the caudate nucleus (REM sleep disorder, gait problems) and the most dorsorostral part of the caudate nucleus has been reported to have a strong reduction in dopamine levels in Parkinson’s disease patients. A recent study thus re-examined the involvement of dopamine deficiencies in the early stage of Parkinson’s disease and found a significant dopaminergic de-innervation of the caudate nucleus in about half of the patients (Pasquini et al., 2019). BCL11A is expressed in human mDA neurons (La Manno et al., 2016), but further studies will be necessary to evaluate which subpopulations express this transcription factor and whether the reduced dopaminergic innervation of the caudate nucleus in Parkinson’s disease may be associated with a specific loss of BCL11A-expressing mDA neurons in patients.

While Bcl11a-mDA neurons are more susceptible to α-synuclein induced degeneration, BCL11A also acts as a neuroprotective factor in this population, since the loss of Bcl11a-mDA neurons is significantly more severe in *Bcl11a* cko^lacZ^ mice than in *Bcl11a-lacZ* mice. A similar phenomenon has been observed for the ALDH1A1 expressing population: inactivation of ALDH1A1 increases the vulnerability of this population to neurodegeneration (Liu et al., 2014). This could be due to the particularly high vulnerability of these populations rather than to a specific function of these factors: any additional insult (i.e. loss of ALDH1A1 or BCL11A function) during development or in the adult brain increases their vulnerability even further. Alternatively, BCL11A could be modulating cell survival more directly, since it has been shown to regulate expression of the anti-apoptotic factor Bcl2 in early B-lymphocytes and inactivation of *Bcl11a* in cortical projections neurons results in increased cell death in addition to deficits in migration and cell fate specification (Yu et al., 2012; Wiegreffe et al., 2015).

## Material and Methods

### Mouse lines

*Bcl11a^lacZ^ mice* (Dias et al., 2016) were kindly provided by Pengtao Liu, School of Biomedical Sciences, The University of Hong Kong, China. *Bcl11a^flox^* mice (Wiegreffe et al., 2015) were kindly provided by Pengtao Liu, School of Biomedical Sciences, The University of Hong Kong, China; and Neal Copeland, Institute for Academic Medicine, Houston Methodist and obtained from Stefan Britsch, University of Ulm. *Bcl11a^CreER^* mice (Pensa et al., under revision) were kindly provided by Walid Khaled, Department of Pharmacology, University of Cambridge. *Bcl11a* cko mice were generated by crossing *Dat^IRES-Cre^* mice (Bäckman et al., 2006) with *Bcl11a^flox^* mice (Genotype: *Dat^RES-Cre/+^, Bcl11a^flox/flox^*). In a subset of *Bcl11a* cko mice, the *Bcl11a^lacZ^* null allele was introduced by crossing *Dat^IRES-Cre^* mice with *Bcl11a^flox/lacZ^ mice* (Genotype: *Dat^IRES-Cre/+^, Bcl11a^flox/lacZ^*). Intersectional fate mapping experiments were performed by crossing *Bcl11a^CreER^* mice with the *Dat^tTA^* (tetracycline trans-activator driven by the *Dat* promoter) mouse line (Chen et al., 2015) and the intersectional reporter mouse line Ai82D (Ai82(TITL-GFP)-D (Madisen et al., 2015)) (Genotype: *Bcl11a^CreER/+^, Dat^tTA/+,^ Ai82D^TITL- GFP/+^*). For viral tracings, *Bcl11a^CreER/+^* or *Bcl11a^CreER/+^, Dat^tTA/+^* (intersectional) were used. Mice were housed in a controlled environment, with 12 hr light/night cycles and ad libitum availability of food and water. Day of vaginal plug was recorded as E0.5. All experiments were performed in strict accordance with the regulations for the welfare of animals issued by the Federal Government of Germany, European Union legislation and the regulations of the University of Bonn. The protocol was approved by the Landesamt für Natur, Umwelt und Verbraucherschutz Nordrhein-Westfalen (Permit Number: 84-02.04.2014.A410, 84-02.04.2016.A238 and 84-12.04.2015.A550).

### Tamoxifen administration

Tamoxifen was administered by oral gavage to pregnant dams at E15.5 (0.05 ml/10g body weight) or to adult mice (0.075 ml/10g body weight) to label *Bcl11a*-expressing neurons. Tamoxifen (Sigma Aldrich) was prepared as a 20 mg/mL solution in corn oil (Sigma Aldrich).

### Tissue processing

Pregnant females were sacrificed by cervical dislocation. Embryos were transferred into in ice cold PBS, decapitated and dissected. P0 pups were decapitated and their brain dissected in ice cold PBS. Heads (E12.5 – E15.5) or brains (P0) were fixed in 4% paraformaldehyde (PFA) overnight at 4°C. Adult mice were anesthetized with an intraperitoneal injection of Ketanest/Rampun or pentobarbital and subsequently perfused transcardially with phosphate buffered saline (PBS), followed by 4% PFA. The tissue was cryopreserved in OCT Tissue Tek (Sakura). Embryonic and P0 tissue was cryosectioned at 14 μm thickness and collected on glass slides, adult brains were cryosectioned at 40 μm thickness and free-floating sections were collected in anti-freeze solution.

For immunofluorescent staining, sections were re-fixed in 4% PFA for 10 min at room temperature (RT) and incubated in 10% NDS in PBS plus 0.2% Triton X-100 (Sigma-Aldrich) (0.2% PBT, used for embryonic and P0 tissue) or in 10% NDS in 0.5% PBT (adult tissue) for 1 hr at RT. Sections were incubated with primary antibody overnight at 4°C in 3% NDS in 0.2% PBT (embryonic and P0 tissue) or in 3% NDS in 0.3% PBT (adult tissue). For staining with the guinea pig anti-BCL11A antibody and in some cases for rabbit anti-TH antibody (**Table 1**) sections were incubated in the primary antibody for 72 hr at RT (guinea pig anti-BCL11A) or at 4°C (rabbit anti-TH antibody). Sections were washed 3 times for 5-10 min in 0.2% PBT (embryonic and P0 tissue) or in 0.3% PBT (adult tissue) and incubated for 2 hr at RT in secondary antibody in 3% NDS in 0.2% PBT (embryonic and P0 tissue) or in 3% NDS in 0.3% PBT (adult tissue). Sections were washed 3 times for 5-10 min in 0.2% PBT (embryonic and P0 tissue) or in 0.3% PBT (adult tissue) and mounted with Aqua Polymount (Polysciences Inc.). For the detection of BCL11A, biotinylated donkey anti-guinea pig antibody followed by Cy3-Streptavidin was used.

Processing of tissue from mice overexpressing *α*-synuclein involved post-fixation with 4% PFA for 24 hr followed by cryopreservation in 30% sucrose solution. Sections were cut at 35 μm thickness using a freezing microtome. Staining of these sections followed previously described protocols (Helwig et al. 2016).

A list of primary and secondary antibodies is provided in **Table 1**.

**Table 1.**
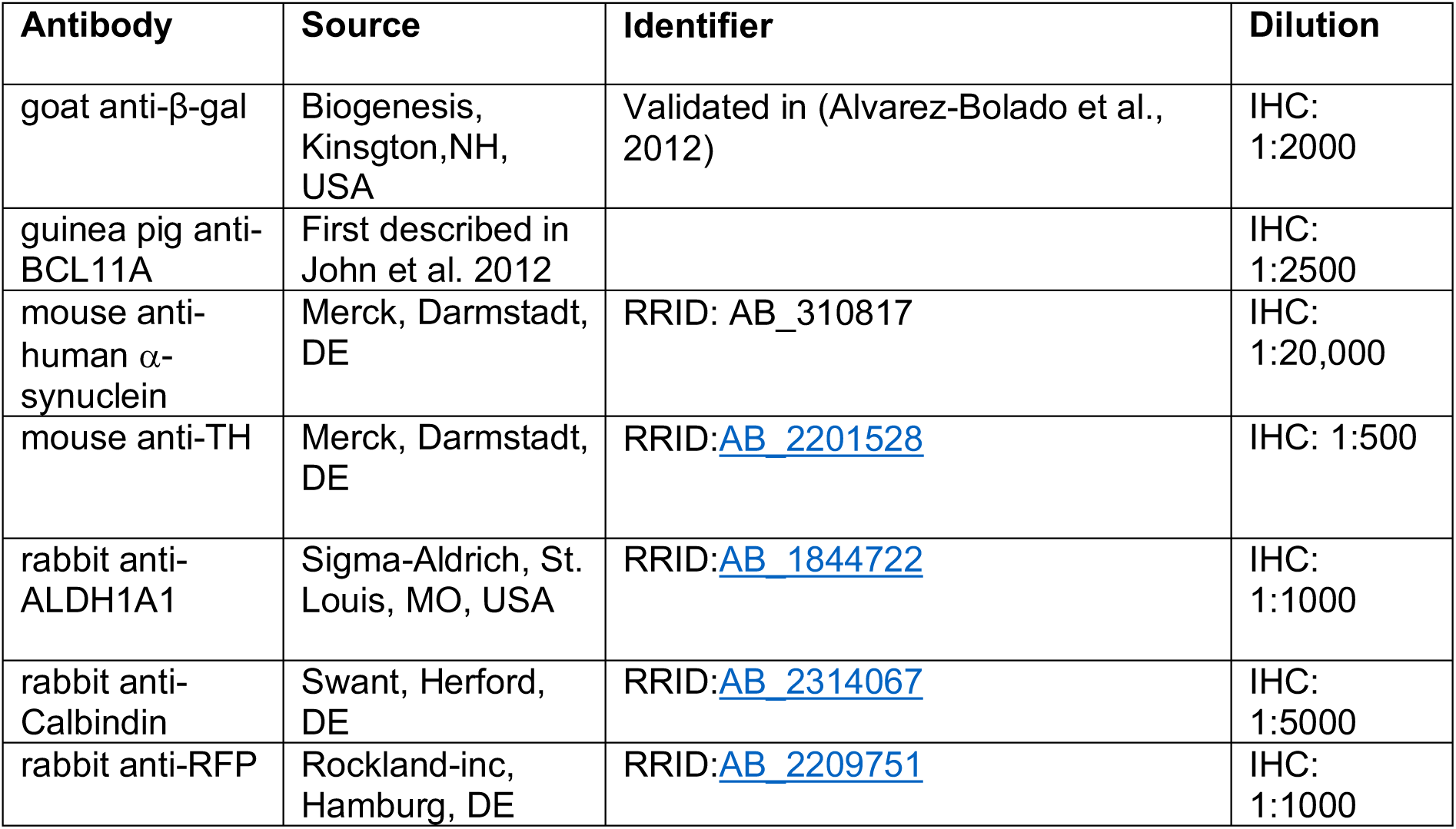
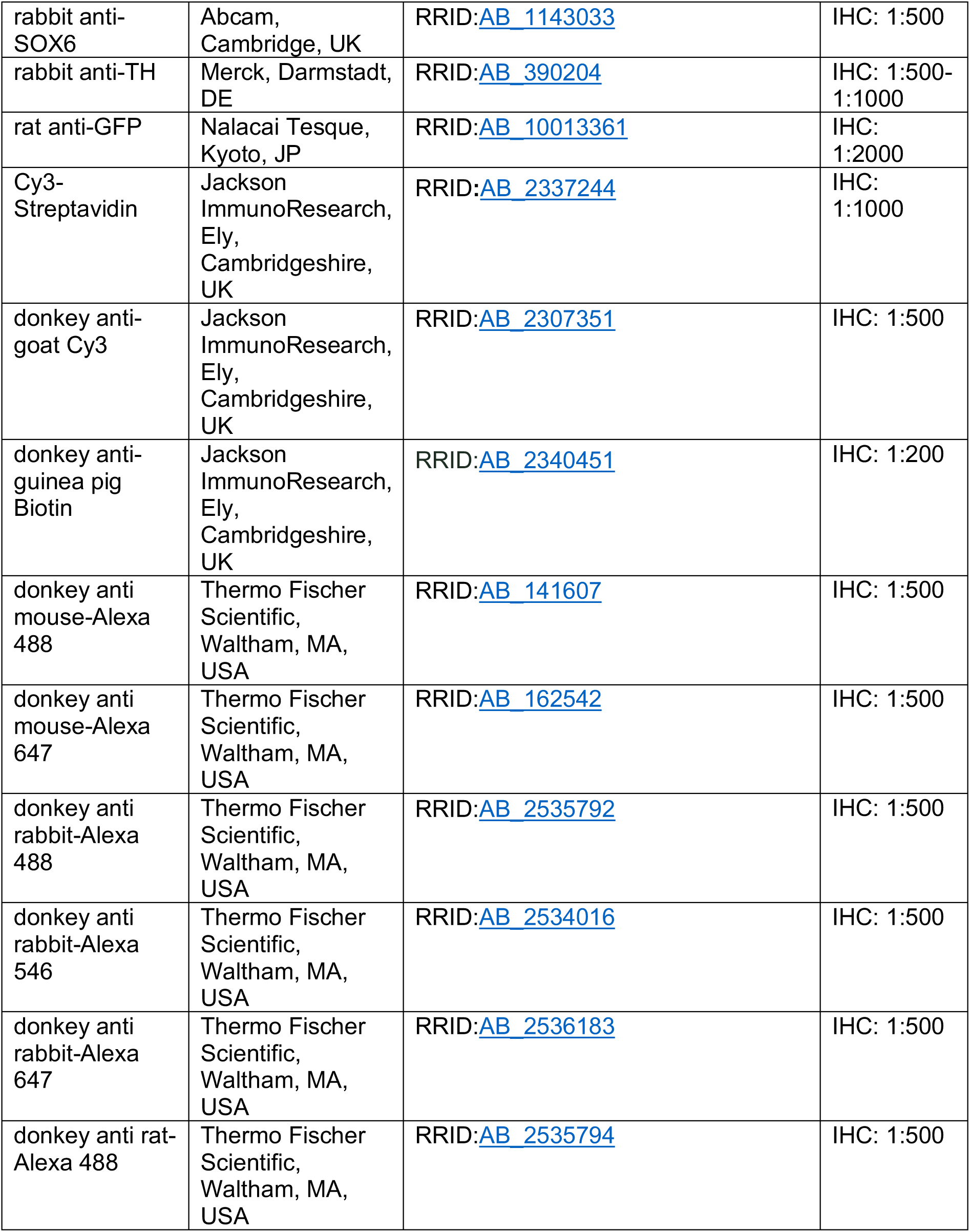

| Antibody | Source | Identifier | Dilution |
| --- | --- | --- | --- |
| goat anti-β-gal | Biogenesis, Kingston, NH, USA | Validated in (Alvarez-Bolado et al., 2012) | IHC: 1:2000 |
| guinea pig anti-BCL11A | First described in John et al. 2012 |  | IHC: 1:2500 |
| mouse anti-human α-synuclein | Merck, Darmstadt, DE | RRID: AB_310817 | IHC: 1:20,000 |
| mouse anti-TH | Merck, Darmstadt, DE | RRID:AB_2201528 | IHC: 1:500 |
| rabbit anti-ALDH1A1 | Sigma-Aldrich, St. Louis, MO, USA | RRID:AB_1844722 | IHC: 1:1000 |
| rabbit anti-Calbindin | Swant, Herford, DE | RRID:AB_2314067 | IHC: 1:5000 |
| rabbit anti-RFP | Rockland-inc, Hamburg, DE | RRID:AB_2209751 | IHC: 1:1000 |
| rabbit anti-SOX6 | Abcam, Cambridge, UK | RRID:AB 1143033 | IHC: 1:500 |
| rabbit anti-TH | Merck, Darmstadt, DE | RRID:AB 390204 | IHC: 1:500-1:1000 |
| rat anti-GFP | Nalacai Tesque, Kyoto, JP | RRID:AB 10013361 | IHC: 1:2000 |
| Cy3-Streptavidin | Jackson ImmunoResearch, Ely, Cambridgeshire, UK | RRID:AB 2337244 | IHC: 1:1000 |
| donkey anti-goat Cy3 | Jackson ImmunoResearch, Ely, Cambridgeshire, UK | RRID:AB 2307351 | IHC: 1:500 |
| donkey anti-guinea pig Biotin | Jackson ImmunoResearch, Ely, Cambridgeshire, UK | RRID:AB 2340451 | IHC: 1:200 |
| donkey anti mouse-Alexa 488 | Thermo Fischer Scientific, Waltham, MA, USA | RRID:AB 141607 | IHC: 1:500 |
| donkey anti mouse-Alexa 647 | Thermo Fischer Scientific, Waltham, MA, USA | RRID:AB 162542 | IHC: 1:500 |
| donkey anti rabbit-Alexa 488 | Thermo Fischer Scientific, Waltham, MA, USA | RRID:AB 2535792 | IHC: 1:500 |
| donkey anti rabbit-Alexa 546 | Thermo Fischer Scientific, Waltham, MA, USA | RRID:AB 2534016 | IHC: 1:500 |
| donkey anti rabbit-Alexa 647 | Thermo Fischer Scientific, Waltham, MA, USA | RRID:AB 2536183 | IHC: 1:500 |
| donkey anti rat-Alexa 488 | Thermo Fischer Scientific, Waltham, MA, USA | RRID:AB 2535794 | IHC: 1:500 |

### RNAScope

RNA in situ hybridization on frozen sections from P0 and adult mice was performed using RNAscope Fluorescent Multiplex Detection Reagents (323110, ACDBio, Newark, CA, USA) according to the instructions provided by the manufacturer for frozen tissue (User Manual: 323100-USM). Hybridized probe was detected with TSA Plus Cyanine 3 (NEL760001KT, Perkin Elmer, Waltham, MA, USA). The probe for Bcl11a was designed by ACDBio (Cat No. 563701-C3). Sections were counterstained with TH and Hoechst, then mounted with Aqua Polymount (Polysciences Inc., Warrington, PA, USA).

### Stereotactic viral vector injection

#### Viral tracing

*Bcl11a^CreER^* and *Bcl11a^CreER^*, *Dat^tTa^* mice (4-20 weeks old) were anesthetized with Fentanyl/Midazolam/Medetomidin and placed into a stereotaxic apparatus. 1 μl of virus solution (AAV1/2-EF1α-pTRE-FLEX-ChETA-eYFP or AAV1/2-EF1α-DIO-ChR2-eYFP (both from UKB viral core facility, Bonn, DE) or AAV1/2-EF1α-DIO-ChR2-mCherry (Addgene, Watertown, MA, USA)) was injected unilaterally into the SNc (from Bregma: anteroposterior (AP), -2.3 mm; mediolateral (ML), -1.1 mm; dorsoventral (DV), -4.1 mm) or into the VTA (from Bregma: AP, -3.44 mm; ML, -0.48 mm; DV, -4.4mm) by. A 34g beveled needle (WPI) and a microinjection pump (WPI) were used to control the injection speed (100 nl/min). After the injection, the syringe was kept in place for 3 min and slowly retracted over 1 min. 8 days after viral injections, Tamoxifen was administered to the mice by oral gavage (0.075 ml/10 body weight) for 3 consecutive days. Two weeks after the last Tamoxifen administration, mice were perfused.

#### Alpha-Synuclein overexpression

Recombinant adeno-associated viral particles (serotype 2 genome and serotype 6 capsid) were used to express human α-synuclein in the mouse substantia nigra. Gene expression was controlled by the human Synapsin 1 promoter and enhanced using a woodchuck hepatitis virus post-transcriptional regulatory element and a polyA signal downstream to the α-synuclein sequence. AAV-vector production, purification, concentration, and titration were performed by Sirion Biotech (Martinsried, Germany). Mice were treated with a single 1.5 µl injection of 4.0×10^12^ genome copies/ml using a stereotaxic frame with a mouse adapter (Stoelting, Wood Dale, IL, USA) under isoflurane anesthesia. Stereotaxic coordinates were 2.3 mm posterior and 1.1 mm lateral to bregma; injection depth was 4.1 mm relative to dura mater. The injection was made at a rate of 0.4µl/min using a Hamilton syringe fitted to a glass capillary. The capillary was left in position for an additional 5 mins before being retracted.

### Image acquisition

Images of fluorescently stained images were acquired at an inverted Zeiss AxioObserver Z1 equipped with structured illumination (ApoTome) and a Zeiss AxioCam MRm (Carl Zeiss, Oberkochen, DE). At 10X (EC PlnN 10x/0.3, Carl Zeiss, Oberkochen, DE) magnification, tile images were acquired with conventional epifluorescence. At 20X (EC PlnN 20x/0.5, Carl Zeiss, Oberkochen, DE), 40X (Pln Apo 40x/1.3 Oil, Carl Zeiss, Oberkochen, DE) and 63X (Pln Apo 63x/1.4 Oil, Carl Zeiss, Oberkochen, DE) magnifications, structured illumination was used to acquire tile images and z-stacks. Some of the images taken with the 20X objective and all the images taken with the 40X and 63X objective are maximum intensity projections of z-stacks. Tile images were stitched with Zen blue software (Zeiss, 2012).

Brightfield images were visualized with a Zeiss Axio Scope.A1 microscope, collected using AxioCam 503 Color and processed with Zen blue software (Zen lite, 2019).

In situ hybridized sections at adult stages were imaged at an inverted Zeiss AxioObserver equipped with a CSU-W1 Confocal scanner unit (50 μm pinhole disk, Yokogawa, Tokyo, JP). At 40X (C-Apochromat, 40x/1.2 water, Zeiss) magnification, tile images and z-stacks were acquired with laser lines 405 nm, 488 nm and 561 nm. Images taken with the 40X objective are maximum intensity projections of z-stacks. Tile images were stitched with VisiView software (Visitron Systems, Puchheim, DE).

### Quantification of cell numbers

#### TH^+^ β-gal^+^ neurons and additional subset markers

The percentage of *Bcl11a*-expressing mDA neurons in SN or VTA at P0 and P30 was determined by quantifying TH^+^ β-gal^+^ neurons at four (P0) or three (P30) rostrocaudal midbrain levels (Franklin and Paxinos, 2007). TH^+^ β-gal^+^ neurons in CLi and RRF in the adult brain were analyzed separately from neurons in the SN or VTA. The number of TH^+^ β-gal^+^ neurons was counted unilaterally in the SN, VTA, RRF and CLi and normalized for the total number of mDA neurons per region separately. The analysis was performed on n=5 *Bcl11a-lacZ* control mice and n=5 *Bcl11a* cko^lacZ^ mice at P0, and on n=6 *Bcl11a-lacZ* (control) mice and n=6 *Bcl11a* cko^lacZ^ mice at P30. The percentage of *Bcl11a*-expressing mDA neurons co-expressing additional subtype markers (SOX6 and CALB1 at P0; CALB1 and ALDH1A1 at P30) in SN or VTA was determined by quantifying TH^+^ β-gal^+^ neurons at four rostrocaudal midbrain levels (Franklin and Paxinos, 2007). The number of TH^+^ β-gal^+^ neurons co-expressing the respective subset marker in these regions was counted unilaterally and the numbers were normalized for the total number of TH^+^ β-gal^+^ mDA neurons in each region (SN or VTA). This analysis was performed for n=5 *Bcl11a-lacZ* mice and n=5 *Bcl11a* cko^lacZ^ mice at P0 and for n=3 *Bcl11a-lacZ* mice and n=3 *Bcl11a* cko^lacZ^ mice at P30.

### Stereology

Analyses were performed on SNc samples by an investigator blinded to the sample codes. Unbiased stereological estimates of the number of nigral neurons were obtained by counting under brightfield or confocal microscopy. Samplings were performed on every fifth section throughout the entire SNc. Delineations were made using a 4x objective, and counting was performed using a 63x Plan-Apo oil objective (Numerical aperture=1.4). A guard zone thickness of 1 µm was set at the top and bottom of each section. Cells were counted using the optical fractionator technique (Stereo Investigator software version 9, MBF Biosciences, Williston, VTA, USA) using a motorized Olympus microscope (IX2 UCB) equipped with an Olympus disk spinning unit (DSU) and a light sensitive EM-CCD camera. Coefficient of error was calculated according to (Gundersen and Jensen, 1987); values <0.10 were accepted.

### Analysis of TH fiber density

40 μm sections of 3 to 6 rostrocaudal levels of the striatum were stained for TH and epifluorescence images were taken using and inverted Z1 Axioobserver microscope with 10X objective. The mean grey value of the TH^+^ striatal projections was calculated in the dorsal striatum, OT and TS (divided into ventral and dorsal TS) with Fiji/ImageJ and normalized for background fluorescence (corpus callosum which is devoid of TH^+^ fibers or neurons). This analysis was performed for =3 control mice and n=3 Bcl11a cko^lacZ^ mice at P30.

### Behavioral tests

Male mice were kept ad libitum in groups of 2-3 animals in a 12-hour day/night rhythm on a normal diet. The animals were transferred to the examination room for experimental purposes and each one was housed in a separate cage. After a one-week acclimatization period in the examination room, mice performed the rotarod test and beam walking assay (both tests in one day) for five consecutive days, followed by the social recognition test and finally by the open-field test.

#### RotaRod

The animals were taken out of the cage and were acclimated to the rotarod apparatus (Ugo Basile, Gemonio, IT; Code no. 47600) in a 5-minute run on a rod rotating at a constant speed of 8 rpm. Afterwards the animals went through three test runs per day, with a break of 30 min between each run. In each test run, the animals had to balance on the rotating rod for 5 min, whereby the torque increased from 4 to 40 rpm within 5 min. The duration that each mouse was able to stay on the rotating rod in each trial was recorded as the latency to fall. The three test runs per day were repeated on 5 consecutive days. During the test, the rod was kept clean and dry by wiping the mouse urine and feces off.

#### Beam Walking Assay

To examine fine motor skills independent of rotational movement (Luong et al., 2011), the beam walking assay (balanced beam test) was used (Carter et al., 1999). Animals had to balance from one platform over a 12 mm wide and 1 m long rod made of synthetic material to another platform that held a box with a food reward. Beams were placed 50 cm above the table. The time taken for the animals to cross the bar was measured. If animals did not reach the safety platform or took longer than 60 s to cross the beam, a maximum of 60 s was assigned. The test was run for a period of 5 consecutive days.

#### Open-field test

In the open-field test, the mice were placed in an open arena (30 x 30 x 30 cm). They were allowed to move freely for 5 minutes. The animals were recorded by Video (EthovisionXT, Noldus, Wageningen, NL) and the running distance, the time spent in the border area, the corners and the centre of the cage as well as crossings from the border to the center area were measured. This test was initially run for a training period of 1 day, then after a 30-day rest period another test run was performed. This test was run twice with a 30-day rest period in between. Since both runs showed comparable results, the results were combined for the final analysis.

### Statistical analysis

Statistical analyses of cell numbers were done with GraphPad Prism (8.0) software using unpaired t-test, Welch’s t-test, one-way ANOVA followed by test for linear trend or one-way ANOVA followed by Tukey’s *post hoc* test for multiple comparisons.

Open field tests were evaluated by one-way analysis followed by Tukey’s post hoc test for multiple comparisons. Differences in the balanced beam test and rotarod were assessed by two-way ANOVA taking time and genotype as numerical and categorical variables.

P values of less than 0.05 were considered statistically significant. Data are reported as mean values ± standard error of the mean (SEM).

## Acknowledgment

This work was supported by the German Research Foundation (BL 767/2–1, BL 767/3–1, BL 767/4–1, 417960915 to Sandra Blaess), the SFB 1089 (to MT and Sandra Blaess), the Maria von Linden-Program (University of Bonn, to Sandra Blaess), and the Ministerium für Kultur und Wissenschaft des Landes Nordrhein-Westfalen (Rückkehrer-Programm to Sandra Blaess). Stefan Britsch has been supported by grants from the DFG (BR 2215/1-1, 2215/1-2). We thank Neal Copeland and Nancy Jenkins for agreeing to provide the Bcl11a floxed mouse line; the UKB microscopy core facility for support with imaging, the UKB virus core for providing AAVs; Milan Pabst and Heinz Beck for support with stereotactical injections, Alesja Dernst, Philipp Grunwald and Julian Wirtz for technical support and Norisa Meli for critical reading of the manuscript.

## Contributions to the paper

**Marianna Tolve**, Data curation, Formal analysis, Validation, Investigation, Visualization, Methodology, Writing—original draft, Writing—review and editing; **Ayse Ulusoy**, Data curation, Formal analysis, Validation, Investigation, Visualization, Methodology, Writing— review and editing; **Khondker Ushna Sameen Islam**; Investigation, Visualization, Methodology, Writing—review and editing; **Gabriela O. Bodea**, Investigation, Visualization, Methodology, Writing—review and editing; **Ece Öztürk, Bianca Broske, Astrid Mentani, Antonia Wagener**, Investigation, Writing—review and editing; **Stefan Britsch**, **Pengtao Liu, Walid Khaled**, **Karen van Loo**, Resources, Writing—review and editing; **Stephan Baader**, Data curation, Formal analysis, Validation, Resources, Methodology, Writing—review and editing; **Donato A. Di Monte**, Data curation, Formal analysis, Validation, Resources, Writing— review and editing; **Sandra Blaess** Conceptualization, Resources, Data curation, Formal analysis, Supervision, Funding acquisition, Validation, Methodology, Project administration, Writing-original draft, Writing—review and editing

---

**Supplemental Figure 1.**
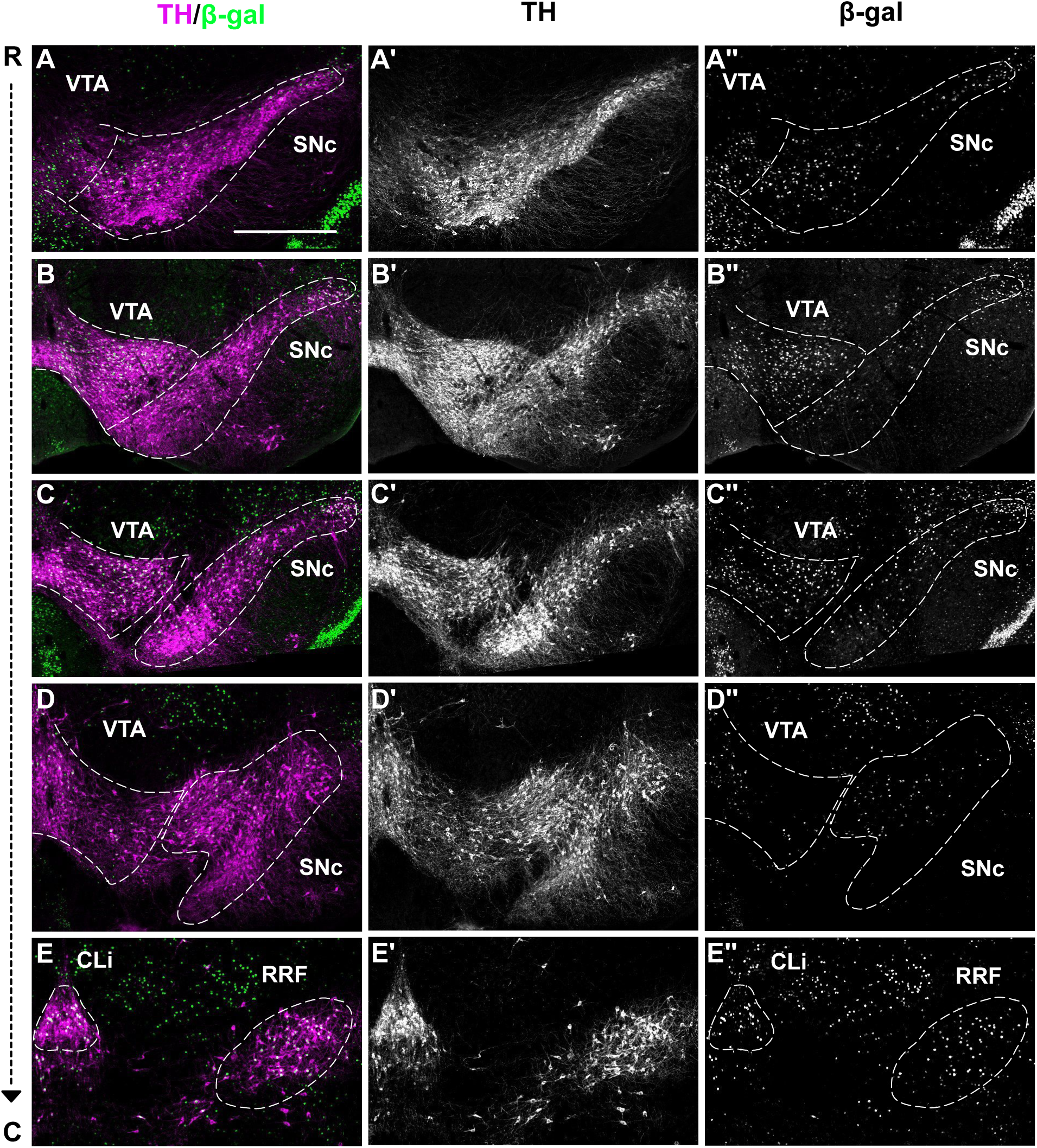
Bcl11a is expressed in a subset of neurons in the SN, VTA, caudal linear nucleus (CLi) and retrorubral field (RRF) of the adult mouse brain. **(A-E′)** Immunofluorescent staining for TH and β-gal on coronal sections at different rostrocaudal levels of P30 *Bcl11a-lacZ* mice. *Bcl11a*-expressing mDA neurons are found throughout the VTA, except for the most medial regions **(A-D′′)**. *Bcl11a*-expressing mDA neurons of the SN are mainly located in the medial SNc (rostral levels) and the dorsal tier of the SNc (more caudal sections) and the SNl **(A-D′′).** *Bcl11a*-expressing mDA neurons are also located in the CLi and RRF **(E-E′′)**. Scale bar: 500 μm **(A-E′′)**.

**Supplemental Figure 2.**
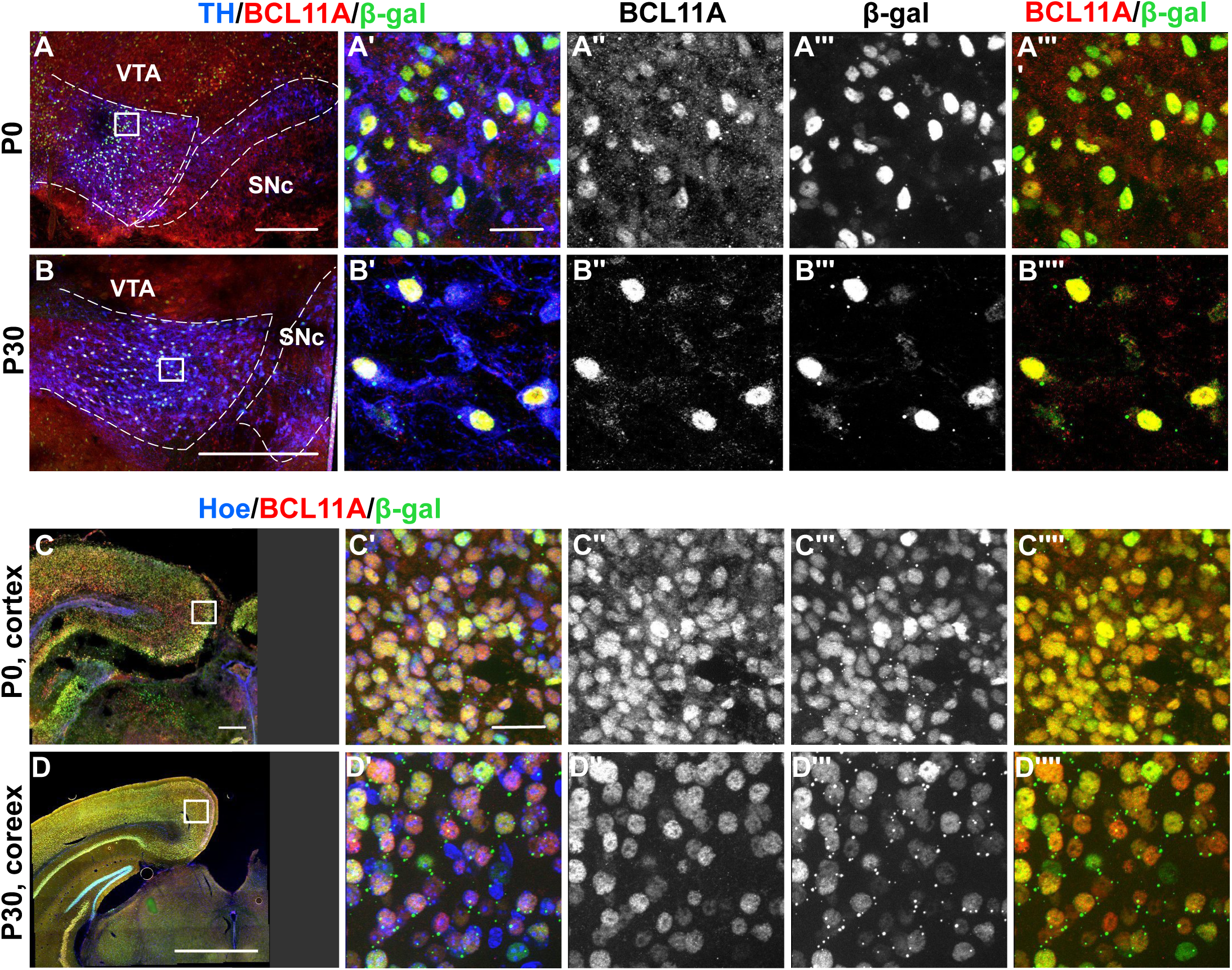
Overlap of BCL11A and β-gal expression in the ventral midbrain and cerebral cortex. **(A-B′′′′)** Triple immunofluorescent staining for TH (blue), BCL11A (red) and β-gal (green) in the ventral midbrain of P0 **(A-A′′′′)** and P30 **(B-B′′′′)** *Bcl11a-lacZ* mice. **(A′-A′′′′, B′-B′′′′)** Higher magnification of the boxed area in A,B. **(C-D’’’’)** Triple immunofluorescent staining for Hoechst (blue), BCL11A (red) and β-gal (green) in cerebral cortex of P0 **(C-C′′′′)** and P30 **(D-D′′′′)** *Bcl11a-lacZ* mice. **(C′-C′′′′, D′-D′′′′)** Higher magnification of the boxed area in C,D. Scale bars: 200 μm **(A,C)**, 500 μm **(B,D)** and 25 μm **(A′-A′′′′,B′-B′′′′,C′-C′′′′,D′-D′′′′)**.

**Supplemental Figure 3.**
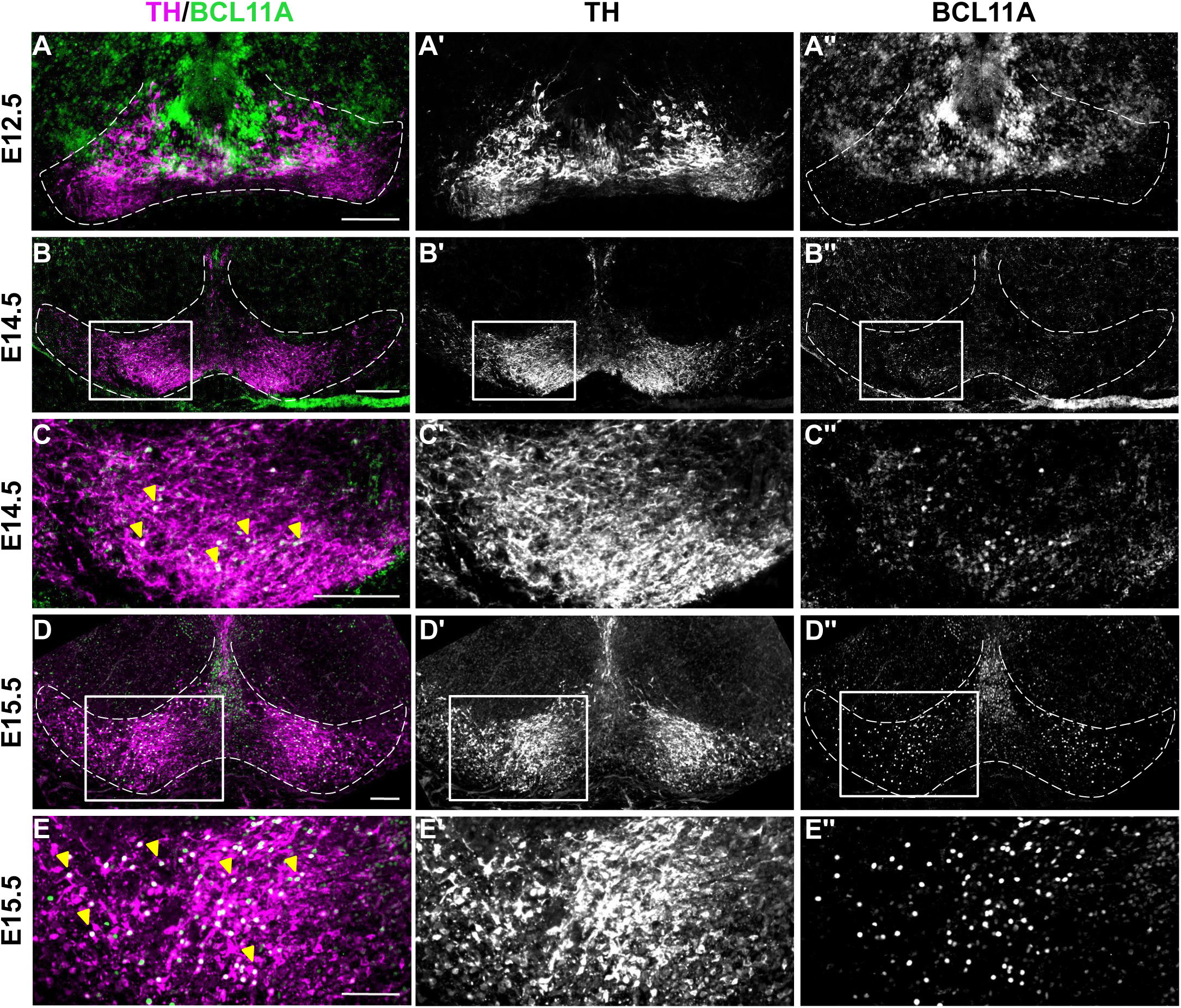
BCL11A expression in the ventral midbrain starts at E12.5. **(A-E′′)** Immunofluorescent staining for TH and BCL11A on E12.5 **(A-A′′)**, E14.5 **(B-C′′)** and E15.5 **(D-E′′)** coronal sections. At E12.5, BCL11A is mainly expressed in differentiated neurons just below the mDA progenitor domain and in a few TH-expressing mDA neurons **(A-A′′)**. At E14.5 and E15.5, BCL11A expression is found in a larger subset of mDA neurons, both in the forming VTA and SN **(B-E′′)**. **(C-C′′, E-E′′)** Higher magnification of the boxed area in B-B′′, D-D′′. Scale bar: 100 μm **(A-E′′)**.

**Supplemental Figure 4.**
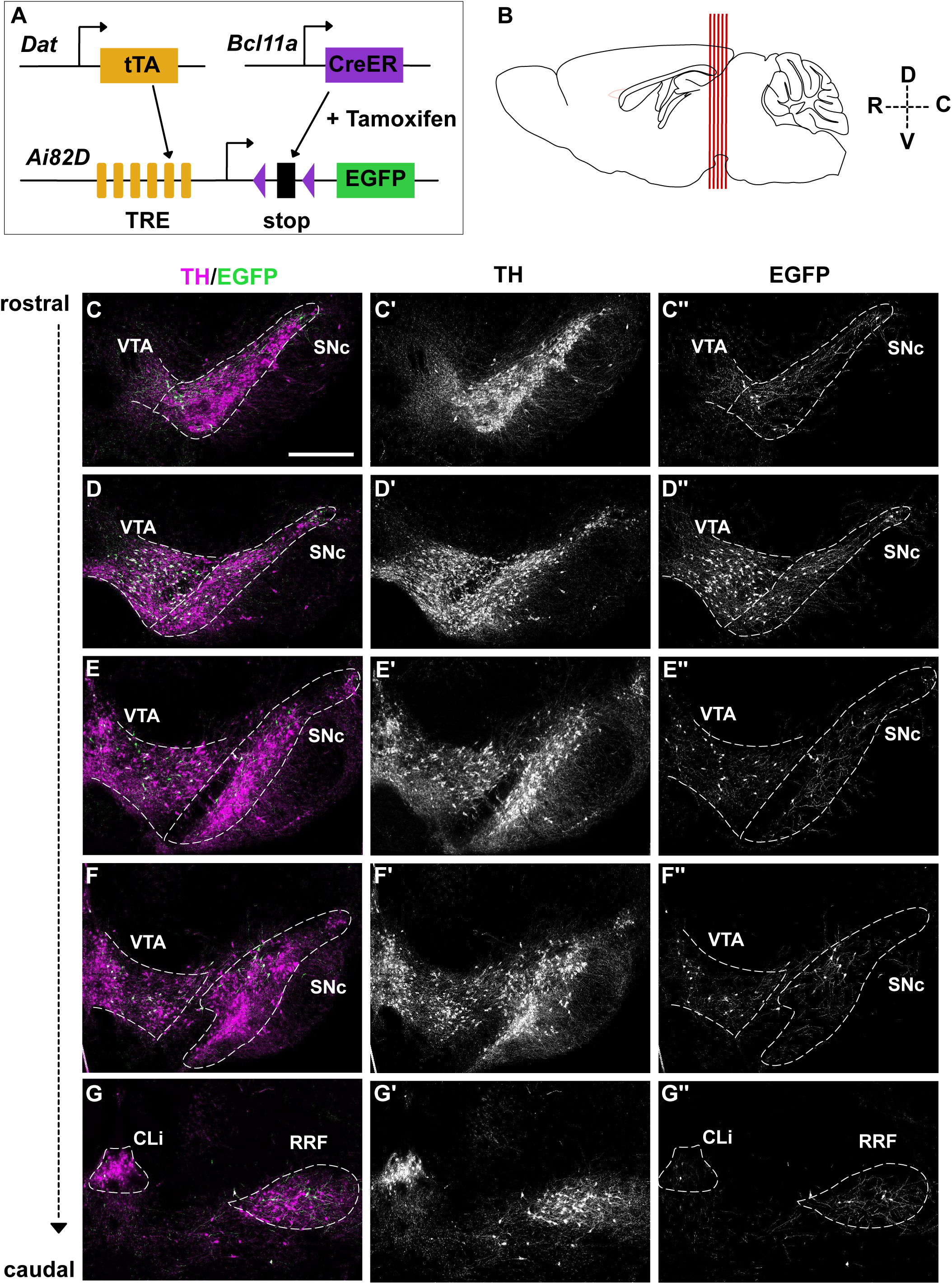
Distribution of recombined neurons in the ventral midbrain using the intersectional fate mapping approach. **(A)** Schematic showing the intersectional fate mapping strategy to label *Bcl11a*-expressing mDA neurons and their projections. **(B)** Schematic of a sagittal section through the adult mouse brain. Red lines indicate the rostrocaudal levels of the coronal sections shown in B-F′′. **(C-G′′)** Immunofluorescent staining for TH and EGFP. Recombined neurons positive for EGFP were found in the SN and VTA **(C-F′′)** as well as in the CLi and RRF **(G-G′′)**. Distribution of recombined cells is comparable to the distribution of β-gal positive cells in *Bcl11a-lacZ* mice (compare with Suppl. Figure 1). n= 8 mice. Scale bar: 500 μm **(B-F′′)**.

**Supplemental Figure 5.**
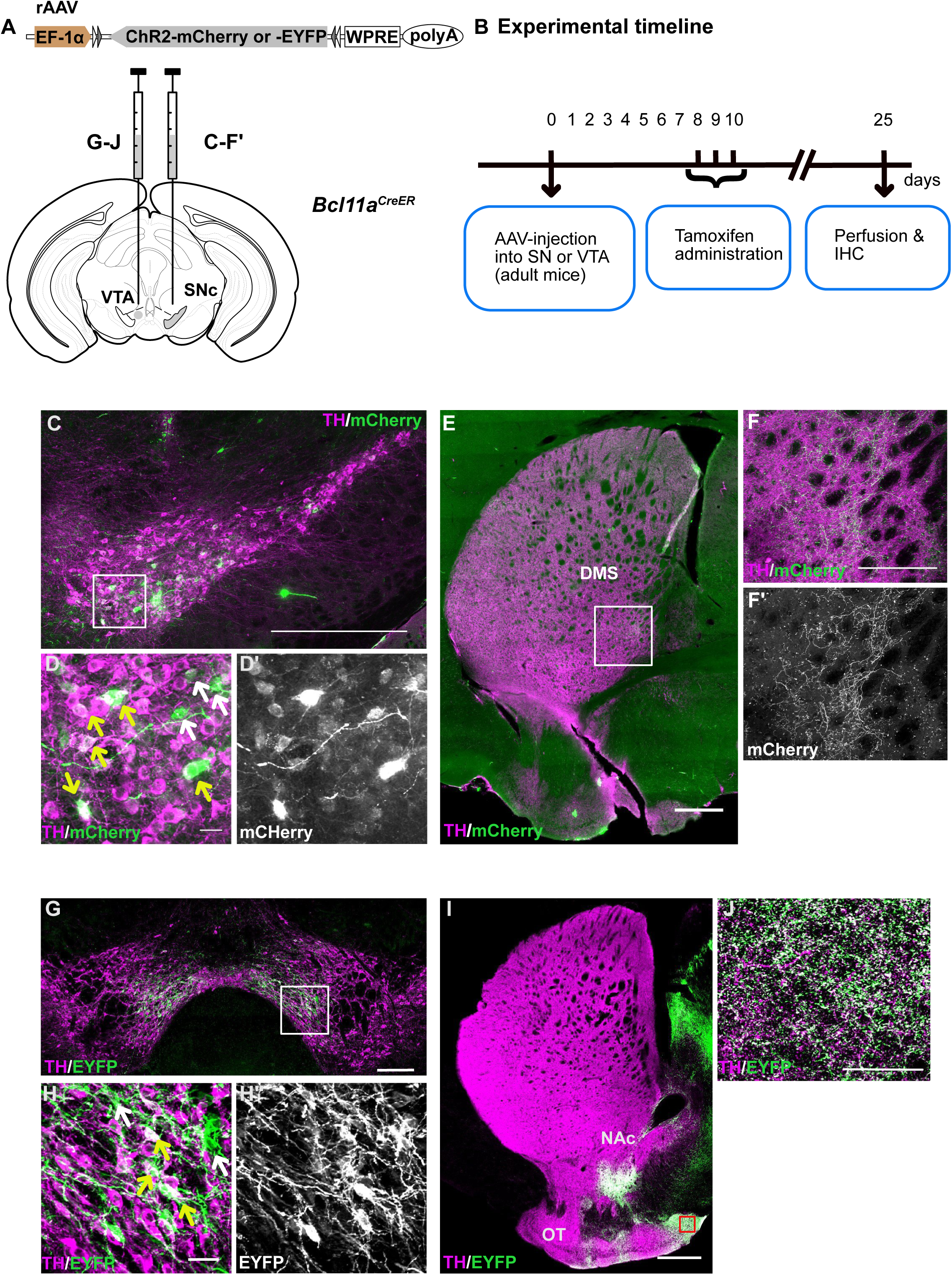
*Bcl11a*-expressing neurons of the VTA and SN show a specific innervation pattern of forebrain targets**. (A)** Schematic showing injection of rAAV:double floxed Channelrhodopsin2 (ChR2)-mCherry or rAAV:double floxed ChR2-EYFP into the SN or VTA of *Bcl11a^CreER^*^/+^ mice. **(B)** Schematic showing the experimental timeline. Tamoxifen administration 8 days after the virus injection leads to sparse expression of the reporter protein (mCherry or EYFP) in *Bcl11a-*expressing neurons. **(C-F′)** Immunofluorescent staining for TH and mCherry in the SN **(C-D′)** and the striatum **(E-F′). (E-F′)** A cluster of mCherry^+^-fibers in the dorsomedial striatum (DMS). **(F-F′)** Higher magnification of the boxed area in E. **(G-J)** Immunofluorescent staining for TH and EYFP in the VTA **(G-H′)** and in the nucleus accumbens (NAc) and the olfactory tubercle (OT) **(I-J). (J)** Higher magnification of the boxed area in I. Yellow arrows indicate TH^+^ reporter protein^+^ neurons, white arrows indicate TH^-^ reporter protein^+^ neurons **(D,H)**. n=7 mice for VTA injections, n=1 mouse for SN injection. Scale bars: 500 μm **(C,E,I)**, 250 μm **(G)** and 25 μm **(F-F′,J)**.

**Supplemental Figure 6.**
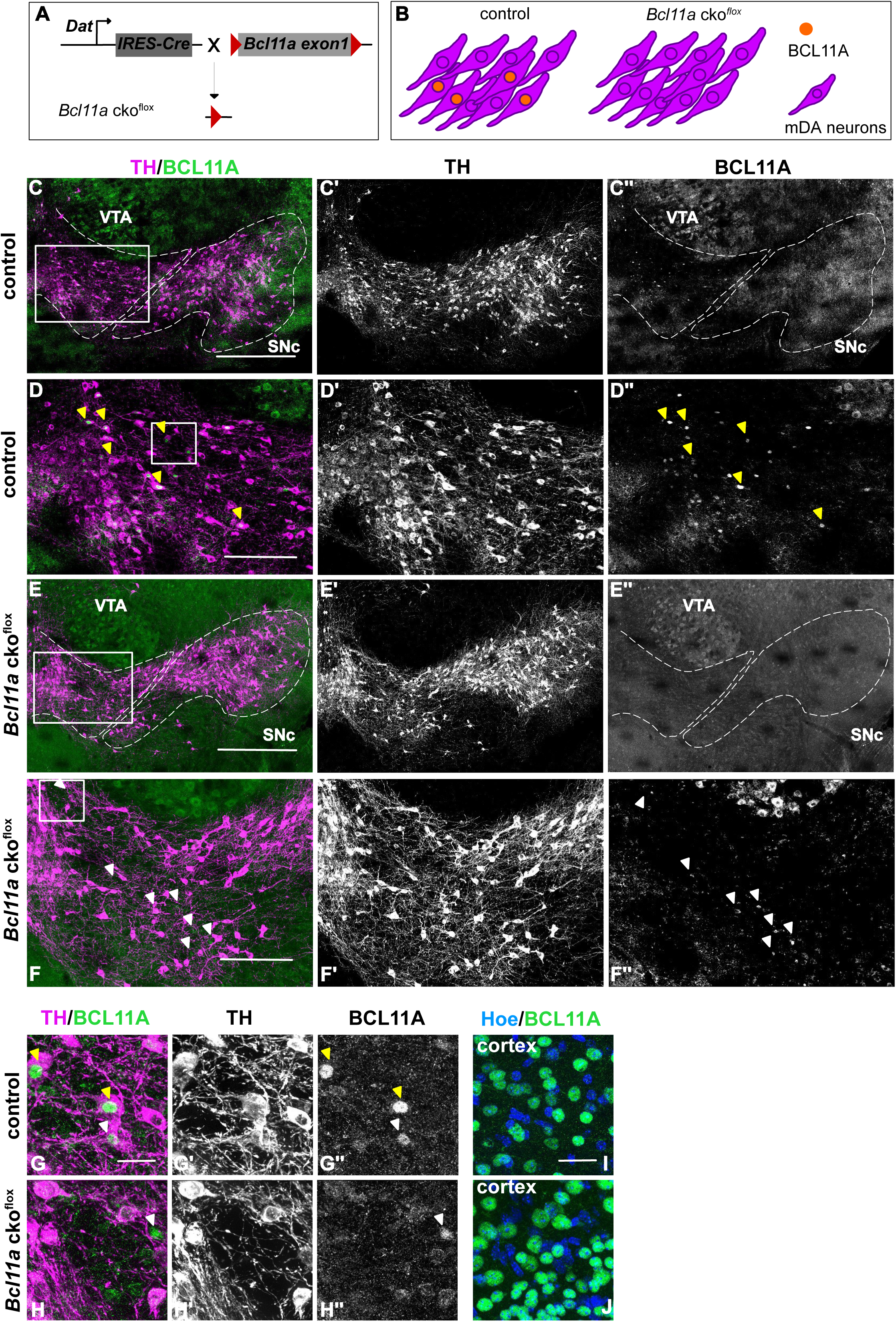
BCL11A expression is absent in mDA neurons of *Bcl11a* cko mice. **(A)** Schematic showing conditional gene inactivation of *Bcl11a* in mDA neurons. Conditional knock-out mice were generated by crossing *Bcl11a^flox/flox^* mice with *Dat^IRES-Cre/+^* mice (*Bcl11a* cko^flox^). **(B)** In *Bcl11a* cko^flox^ mice, BCL11A expression is absent in mDA neurons. **(C-H′′)** Immunofluorescent staining for BCL11A and TH on control (C-D’’) and *Bcl11a* cko^flox^ (E-F**′′**) coronal sections of the adult brain. BCL11A expression is present in mDA neurons of control mice **(C-D′′’, G,** yellow arrowheads), but it is absent in mDA neurons of *Bcl11a* cko^flox^ mice **(E-F′′, H)**. BCL11A is still expressed in non-dopaminergic neurons of the *Bcl11a* cko^flox^ mice midbrain **(E-F′′, G,** white arrowheads). **(D-D′′,F-F′′)** Higher magnification of the boxed area in C,E. **(G-G′′,H-H′′)** Higher magnification of the boxed area in D,F. Yellow arrowheads indicate cells that are double positive for BCL11A and TH **(D,D′′,G,G′′)**, white arrowheads indicate cells that are BCL11A positive but TH negative **(F,F′′,G,G′′,H,H′′)**. **(I-J)** Immunofluorescent staining for BCL11A and Hoechst in control **(I)** and *Bcl11a* cko^flox^ **(J)** cerebral cortex. As expected, BCL11A is still expressed in cerebral cortex neurons of *Bcl11a* cko^flox^ mice **(J)**. Scale bars: 500 μm **(C-C′′,E-E′′**), 250 μm **(D-D′′,F-F′′)** and 25 μm **(G-J)**.

**Supplemental Figure 7.**
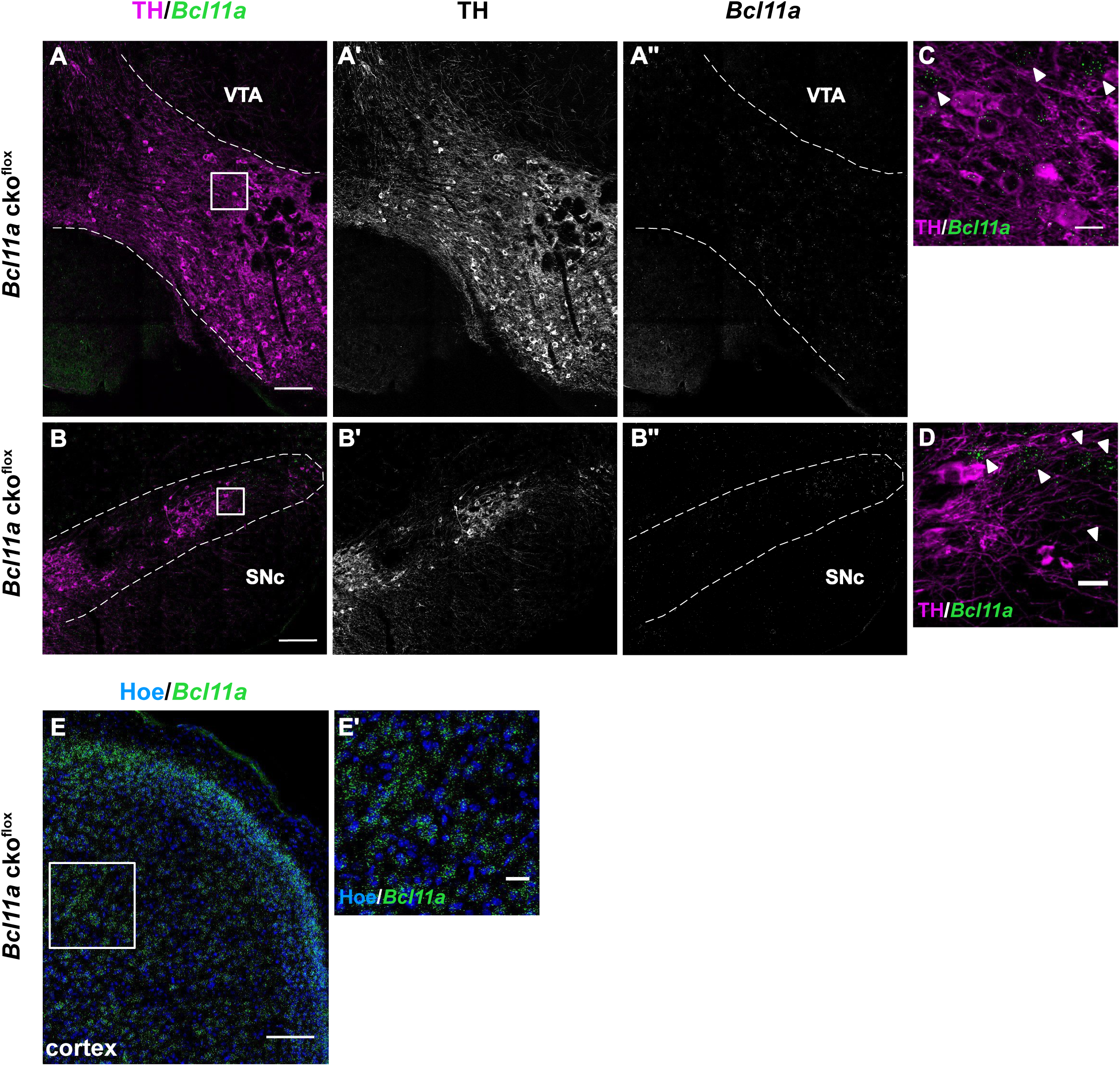
*Bcl11a* mRNA is no longer expressed in mDA neurons of *Bcl11a* cko mice. **(A-D)** Immunofluorescent staining for TH and RNAscope for *Bcl11a* on *Bcl11a* cko^flox^ coronal sections of the adult brain. *Bcl11a* mRNA expression is still expressed in non-dopaminergic neurons of the *Bcl11a* cko^flox^ mice VTA **(A-A′′,C)** and SN **(B-B′′,D)**. **(C-D)** Higher magnification of the boxed area in A,B. White arrowheads indicate cells that are expressing *Bcl11a* mRNA but that are TH negative. **(E,E′)** Immunofluorescent staining for Hoechst and RNAscope for *Bcl11a* in *Bcl11a* cko^flox^ cerebral cortex. As expected, cerebral cortex neurons of *Bcl11a* cko^flox^ mice are still expressing *Bcl11a* mRNA. **(E′)** Higher magnification of the boxed area in E. Scale bars: 250 μm **(A-B′′)**, 200 μm **(E)** and 25 μm **(C,D,E′)**.

**Supplemental Figure 8.**
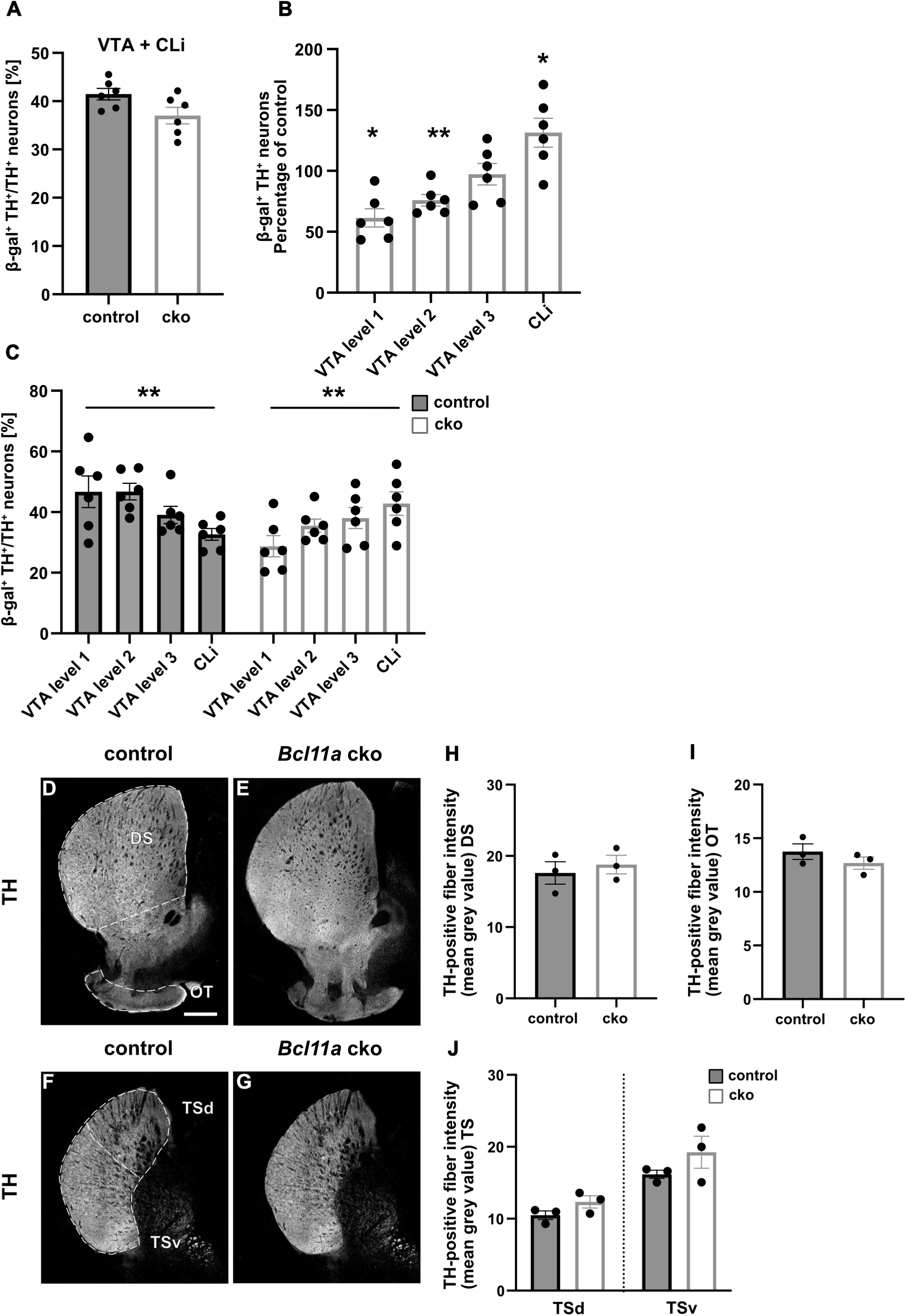
*Bcl11a* inactivation does affect rostro-caudal position of Bcl11a-mDA neurons but not the targeting of projections to areas innervated by *Bcl11a*-expressing mDA neurons. **(A)** Quantification combining three rostrocaudal VTA levels and the CLi. No significant difference in the percentage of β-gal positive TH neurons could be detected between *Bcl11a* cko^lacZ^ and *Bcl11-lacZ* animals. Statistical significance was determined by unpaired t-test. **(B)** Percentage of β-gal positive TH neurons normalized to control three rostrocaudal VTA levels and the CLi (the CLi data are included for easier comparison the same values are also shown in Figure 5F). The percentage of β-gal positive TH neurons at the two rostral VTA levels (level 1 and 2) was significantly decreased in *Bcl11a* cko^lacZ^ compared to control mice, while there was a significant increase in the CLi. Statistical significance was determined by Welch’s t-test. **(C)** In control animals, there is a systematic decrease in β-gal positive TH neurons from rostral-to-caudal (level1 to CLi), while there is a systematic increase in the percentage of β-gal positive TH neurons from rostral-to-caudal in *Bcl11a* cko^lacZ^ mice Statistical significance was determined by one-way ANOVA followed by a test for linear trend. **(D-G)** Immunofluorescent staining for TH on striatal sections of control **(D,F)** and *Bcl11a* cko^lacZ^ **(E,G)** mice. **(H-J)** Quantification of TH-positive fiber density in dorsal striatum (DS) **(H)**, olfactory tubercle (OT) **(I)** and ventral (v) and dorsal (d) TS **(J)** of control (n=3 mice) and *Bcl11a* cko mice (n=3 mice) shows no difference in the density of TH innervation between the two groups. Statistical significance was determined by unpaired t-test. Error bars indicate mean +/- SEM. Scale bar: 500 μm **(A-D)**.

